# Evidence for coordinated evolution at amino acid sites of SARS-CoV-2 spike

**DOI:** 10.1101/2022.08.30.505885

**Authors:** A.D. Neverov, G.G. Fedonin, A.V. Popova, D.I. Bykova, G.A. Bazykin

## Abstract

It is currently unclear why SARS-Cov-2 has adapted in a stepwise manner, with multiple beneficial mutations accumulating in a rapid succession at the origins of VOCs. Here, we searched for coordinated evolution of amino acid sites in the spike protein of SARS-Cov-2. We searched for concordantly evolving site pairs (CSP) for which changes at one site were rapidly followed by changes at the other site in the same lineage. We detected 46 sites which formed 45 CSP. Sites in CSP were closer to each other in the protein structure than random pairs, indicating that concordant evolution has a functional basis. Notably, site pairs carrying lineage defining mutations of the four VOCs that circulated before May 2021 are enriched in CSP, indicating that the origin of these VOCs could have been facilitated by positive epistasis. Additionally, we detected four discordantly evolving pairs of sites where mutations at one site unexpectedly rarely occurred on the background of a specific allele at another site, namely on the wild-type D at site 614 (for two pairs) or at derived Y in the site 501 (for two other pairs). Our findings hint that positive epistasis between accumulating mutations could have delayed the assembly of advantageous combinations of mutations comprising at least some of the VOCs.

## Introduction

Evolution of SARS-CoV-2 in human hosts before November 2020 was largely neutral, with little evidence for emergence of novel adaptation with the exception of fixation of D614G in the Spike protein (Dearlove et al., 2020; MacLean et al., 2021). However, since the end of 2020, evidence for adaptive viral evolution has started to accumulate, suggesting a change in the mode of evolution (D. P. Martin et al., 2021; Rochman et al., 2021b). The subsequent pandemic was characterized by emergence of multiple concurrently circulating lineages with increased fitness compared to the ancestral variant, including Alpha (B.1.1.7), Beta (B.1.351), Gamma (P.1) and Delta (B.1.617.2). These lineages are typically characterized by high divergence, compared to cocirculating strains; divergence is often particularly pronounced at nonsynonymous sites, suggesting positive selection at origin of these variants. Some of these sites are evident of a change in selection regime at the origin of VOCs. For example, out of the 34 lineage-defining amino acid changes in the S-protein at the origin of the Omicron BA.1 sublineage (Emma B. Hodcroft., 2021), eleven were characterized by strong purifying selection against changes of ancestral amino acids (Martin et al., 2022), suggesting that Omicron lineage-defining mutations at these sites were previously individually deleterious. In turn, the obvious high fitness of Omicron suggested that the origin of this variant has been characterized by a change in the selection regime at least at these sites. The reasons for this change are unclear. Several non-exclusive explanations were proposed, including a distinct mode of evolution at variant origin, e.g., in an immunosuppressed individual (Corey et al., 2021; Kai Kupferschmidt, 2021) or a different host species (Wei et al., 2021); and/or cascades of substitutions at positively epistatically interacting sites (Moulana et al., 2022). Here, we focus on the latter possibility.

Several previous studies have attempted to infer possible epistatic interactions between sites of SARS-CoV-2 genome from sequence data or experimentally. In an early study, Zeng et. al. used direct coupling analysis (DCA) to search for epistasis in SARS-CoV-2 genome and reported several pairs of putatively interacting sites (Zeng et al., 2020). No pairwise interactions between sites of the S-protein were identified. For DCA to accurately detect interacting sites, the analyzed sequences need to be highly divergent (Bisardi et al., 2022). SARS-CoV-2 has a recent common ancestor, and divergence of its lineages is relatively low, limiting the application of DCA for this virus. Rodriguez-Rivas et al. applied DCA to homologous protein sequences from genomes of other coronaviruses and successfully predicted variability of SARS-CoV-2 protein sites, thus showing that knowledge of covariation between sites in related viruses is relevant for predicting evolution of new pathogens (Rodriguez-Rivas et al., 2022). In the study of Rochman et al., a method based on counting of mutations on the phylogeny was used to look for strongly associated mutation pairs (Rochman et al., 2021b, 2020). They found intra- and intergenic epistasis between positively selected mutations in the nuclear localization signal (NLS) of the N-protein and RBD in the S-protein. In RBD, many of the detected epistatically interacting mutations were among the lineage signature mutations. Another proposed approach to study epistatic interaction was to estimate the fitness effects of mutations that arose on the different backgrounds relative to their effects on the wild type background (Rochman et al., 2021a). Using molecular dynamics, Rochman et. al. estimated effects of all individual non-synonymous mutations in the S-protein RBM on binding with host ACE2 receptor and on binding with neutralizing Ab (NAb). Effects of each mutation were estimated for the Wuhan ancestral background, Delta (452R, 478K), Gamma variants (417T, 484K, 501Y) and Omicron (339D, 371L, 373P, 375F, 417N, 440K, 446S, 477N, 478K, 484A, 493R, 496S, 498R, 501Y, 505H). On average, the epistatic effects of mutations weakly stabilized NAb binding for Delta and destabilized it for Gamma and Omicron variants relative to the ancestral background. The authors concluded that the Gamma and Omicron variants had a higher potential for emergence of immune escape mutations than Delta or Wuhan variants.

For some site pairs, epistasis had been demonstrated experimentally. For example, the Q498R mutation alone affected the affinity of Spike to ACE2 only slightly (Zahradník et al., 2021), but on the background of N501Y, the affinity of binding increased by a factor of 4 to 25 (Starr et al., 2022; Zahradník et al., 2021), with both mutations together increasing the affinity by up to 387-fold compared to the wild type (Starr et al., 2022). The very strong binding provided by the double mutant allows accumulation, at Omicron origin, of multiple immune escape mutations at other sites which by themselves destabilize ACE2 binding (Starr et al., 2022).

Here, we study the mutual distribution of spike mutations in SARS-CoV-2 phylogeny to infer the pairs of sites with evidence for concordant and discordant evolution, as manifested by the propensity of substitutions at these sites to occur rapidly one after the other (for concordant evolution), or to avoid each other (for discordant evolution). We detect 46 concordantly evolved sites combined into 13 coevolving clusters, and six discordantly evolved sites. Many of the concordantly evolved sites carry the characteristic mutations of VOC lineages, strongly suggesting the role of positive epistasis in VOC origin.

## Results

### Detecting interdependently evolving pairs of sites

To find coevolving site pairs, we modified our previously developed phylogenetic approach (Kryazhimskiy et al., 2011; Neverov et al., 2021, 2015) to improve the accuracy of detecting concordantly evolving site pairs (see Methods). Similarly to our previous work, as a measure of concordance of evolution at two sites, we used the epistatic statistics calculated as the weighted sum of consecutive pairs of mutations at these two sites on the phylogeny, where each mutation pair was taken with exponential penalty for the waiting time for the later mutation (Kryazhimskiy et al., 2011).

We need to introduce some definitions for further explanation. Hereafter, unless specified otherwise, we use the term ‘mutation’ for defining a triple of a site, ancestral and derived amino acids identifiers. Using ancestral state reconstruction, we are able to infer the order in which two specific mutations occurred in an evolving lineage. For a pair of consecutive mutations at two sites, we call the mutation that occurs first a leading mutation, and the mutation that follows it, a trailing mutation. For an ordered pair of sites, we call the first site in a pair the background site, and the second site in the pair, the foreground site. The epistatic statistic for an ordered pair of sites summarizes the weights of consecutive pairs of mutations at these sites, such that mutations at the background site are leading and mutations at the foreground site are trailing. The epistatic statistics for an unordered pair of sites is a sum of statistics of the two corresponding ordered pairs (Neverov et al., 2021).

We introduced two significant changes to the original method (Kryazhimskiy et al., 2011; Neverov et al., 2021) which improved the power to infer epistasis (“revised method”, see next section). First, as in (Neverov et al., 2015), we modified the null model used to calculate the significance of the epistatic statistics in permutations. While previously (Kryazhimskiy et al., 2011; Neverov et al., 2021) we permuted the positions of mutations on the tree branches at each site independently of other sites, here, we fixed the positions of mutations for the background site and permuted just the positions of mutations at the foreground site. This change allowed us to account for the possible effects of leading mutations on the topology of the phylogenetic tree; e.g., an advantageous leading mutation could give rise to a prolific clade (Neher, 2013) which in turn would carry a large number of trailing mutations, artefactually inflating the epistatic statistic for this pair of sites even in the absence of epistatic interactions.

Second, while our previous work (Kryazhimskiy et al., 2011; Neverov et al., 2021) treated all substitutions at a site equally, we now distinguished between substitutions into different amino acids. Therefore, the revised epistatic statistic accounts for the preference of a specific mutation at the foreground site to follow a specific mutation at the background site. For this, in calculation of the epistatic statistic, we now additionally scored each pair of consecutive mutations by the fraction of times that the specific type of mutation at the foreground site followed the specific type of mutations at the background site, among all occurrences of this type of mutations at the foreground site. Therefore, extra weight was given to those mutation types that became more frequent on a specific background.

### Estimating the power of the method to detect epistasis

To demonstrate that our revised method improves inference of positive and negative epistasis, we used MimicrEE2 (Vlachos and Kofler, 2018) to simulate clonal evolution of linked sites. We simulated two modes of evolution: (i) under positive and negative selection without epistatic interactions (“multiplicative mode”), and (ii) under epistatic selection (“epistatic mode”).

Specifically, we simulated independent forward-time evolution of a population of 50000 genotypes consisting of 100 biallelic sites. For the multiplicative mode, at the start of the simulations, twenty sites of the gene carried the disfavored allele, and were therefore under positive selection; and twenty sites carried the favored allele, and were therefore under negative selection. For the epistatic mode, twenty sites constituted ten site pairs such that the sites within each pair evolved under positive epistasis; and another twenty sites constituted ten site pairs such that the sites within each pair evolved under negative epistasis. Under each mode, the remaining 60 sites evolved neutrally (see Methods).

We used simulations to estimate how the changes to the method for inference of epistasis introduced in this work impacted method accuracy. For this, using simulated datasets, we compared the power of the four variants of our method, corresponding to the presence or absence of the two modifications introduced in the previous section (accounting for amino acid identities and unlinking the distributions of mutations on the tree branches for background and foreground for the null model).

To compare the specificity of the four variants of the method, we used the multiplicative mode of simulation (i.e., the absence of the epistasis), and asked how frequently concordant or discordant pairs were inferred under each model. Since there was no epistasis in the simulation, each such pair was spurious, and the best method would be the one with fewest such pairs. For each method, we counted the number of spuriously inferred concordant and discordant pairs at the lowest p-value threshold in our simulation trials (10^-4^). There were 24 concordant pairs and 16 discordant pairs in the method of (Neverov et al., 2021), but just 2 concordant pairs and 2 discordant pairs in the revised method, indicating that the modifications introduced here helped improve the specificity of our approach. We used the 10% FDR level for this analysis and for all analyses below. For the FDR 10%, no concordant or discordant pairs were inferred in this dataset by the revised method (tables S1 and S2, figures S1 and S2)..

**Table 1.** Coevolving sites of the SARS-Cov-2 S-protein with FDR less than 10% for both reconstructed phylogenies (see Text). The following characteristics are shown: coordinates on the S-protein sequence, nominal p-values, the value of the epistatic statistics, the total number of consecutive mutation pairs for the two corresponding ordered site pairs, numbers of mutations in consecutive pairs at site1 and site2, total numbers of mutations at site1 and site2, and the distance in the protein structure (PDB ID: 7JJJ).

| site1 | site2 | p-value | epistatic statistics | #consec. pairs of mutations | #mutations in consec. pairs in site1 | #mutations in consec. pairs in site2 | #mutations in site1 | #mutations in site2 | physical distance, Å |
| --- | --- | --- | --- | --- | --- | --- | --- | --- | --- |
| 13 | 152 | <2e-5 | 2.864 | 5 | 5 | 4 | 5 | 16 |  |
| 20 | 417 | 2.2e-4 | 1.348 | 4 | 3 | 2 | 8 | 7 | 47.83 |
| 26 | 190 | 1.8e-4 | 1.029 | 4 | 4 | 3 | 12 | 3 | 18.4 |
| 63 | 213 | 4e-5 | 0.681 | 1 | 1 | 1 | 2 | 3 | 13.51 |
| 69 | 70 | <2e-5 | 1.125 | 2 | 2 | 2 | 5 | 4 | 1.3 |
| 70 | 144 | <2e-5 | 1.301 | 3 | 3 | 3 | 4 | 5 | 14.03 |
| 76 | 490 | 2.6e-4 | 0.22 | 1 | 1 | 1 | 3 | 3 | 45.18 |
| 189 | 360 | 1.6e-4 | 0.544 | 2 | 1 | 2 | 3 | 3 | 29.89 |
| 190 | 417 | 1e-4 | 1.272 | 3 | 2 | 2 | 3 | 7 | 39.93 |
| 259 | 261 | <2e-5 | 1.35 | 1.5 | 2 | 1 | 4 | 2 | 3.81 |
| 356 | 360 | <2e-5 | 1.283 | 2.5 | 2 | 3 | 2 | 3 | 10.12 |
| 359 | 360 | <2e-5 | 0.976 | 1.5 | 1 | 2 | 2 | 3 | 1.34 |
| 439 | 441 | <2e-5 | 2.097 | 4 | 3 | 4 | 9 | 8 | 3.5 |
| 440 | 441 | 1.4e-4 | 1.505 | 3 | 3 | 3 | 12 | 8 | 1.33 |
| 440 | 442 | <2e-5 | 2.476 | 3 | 2 | 3 | 12 | 6 | 3.92 |
| 440 | 444 | <2e-5 | 2.515 | 3.5 | 2 | 5 | 12 | 7 | 5.6 |
| 441 | 442 | <2e-5 | 2.244 | 4 | 4 | 4 | 8 | 6 | 1.33 |
| 441 | 443 | 2e-5 | 1.281 | 5 | 4 | 2 | 8 | 2 | 3.23 |
| 441 | 444 | <2e-5 | 3.492 | 5.5 | 4 | 4 | 8 | 7 | 2.6 |
| 442 | 443 | 6e-5 | 1.301 | 4 | 4 | 2 | 6 | 2 | 1.32 |
| 442 | 444 | <2e-5 | 3.013 | 6 | 4 | 4 | 6 | 7 | 4.05 |
| 443 | 444 | 8e-5 | 1.043 | 2 | 2 | 2 | 2 | 7 | 1.32 |
| 501 | 1118 | 1.4e-4 | 2.737 | 16 | 13 | 12 | 40 | 22 | 131.71 |
| 681 | 716 | <2e-5 | 3.905 | 16.5 | 12 | 16 | 59 | 21 |  |
| 716 | 982 | 2e-5 | 2.001 | 15 | 9 | 11 | 21 | 15 | 81.66 |
| 716 | 1118 | 4e-5 | 2.382 | 13 | 8 | 12 | 21 | 22 | 22.29 |
| 859 | 950 | <2e-5 | 2.219 | 5.5 | 4 | 5 | 11 | 9 | 15.17 |
| 982 | 1118 | 1.4e-4 | 1.637 | 15 | 11 | 8 | 15 | 22 | 94.63 |

To study the accuracy of the four variants of the method, we used simulations with epistasis. The revised method detected all 10 positively epistatic site pairs as concordantly evolving; additionally, it spuriously detected 5 other site pairs as concordantly evolving (tables S3, S4). The revised method also detected 7 out of 10 negatively epistatic site pairs as discordantly evolving, and spuriously detected 4 other site pairs as discordantly evolving (tables S5, S6). The three other detection models were less accurate: the number of false predictions was greater than the number of true predictions for positive epistatic pairs for all other detection methods, and for negative epistatic pairs, for two out of three methods (tables S3, S5). Therefore, the revised method was the method of choice for subsequent analyses.

### Phylogenetic analysis of SARS-CoV-2 spike

To obtain a phylogeny representative of SARS-CoV-2 diversity, we downloaded 3,299,439 complete genome sequences of SARS-CoV-2 aligned to the WIV04 reference genome from the GISAID EpiCov™ database on 07.09.2021. We ignored insertions and deletions relative to the reference sequence and removed sequences with inframe stop codons in the spike protein. We then clustered the remaining sequences by pairwise distances between S-protein subsequences, allowing up to three mutations in the S-protein within a cluster, which resulted in 7,348 clusters. For each cluster, we selected one representative sequence of the best quality with the earliest date of sampling. The median date of representative sequences was February 10, 2021. Therefore, the dataset covered approximately equally both characteristic periods of SARS-CoV-2 evolution: the neutral period between Jan-Nov 2020, and the period of antigenic drift between Dec 2020 – May 2021 (MacLean et al., 2021; Martin et al., 2021). We classified representative sequences according to pangolin lineages. Most sequences (5,721) were of the B.1.* sublineages. The representative sequences included some from the variants of concern (VOCs) Alpha (B.1.1.7+Q.*, 951 sequences), Beta (B.1.351.*, 192 sequences), Delta (B.1.617.2+AY.*, 24 sequences) and Gamma (P.1.*, 100 sequences). The phylogeny of representative sequences was reconstructed using IQ-TREE (Minh et al., 2020). The tree was rooted by the outgroup USA-WA1/2020 (EPI_ISL_404895) that matched the sequence of the putative SARS-CoV-2 progenitor (Bloom, 2021; Kumar et al., 2021). The ancestral sequences at internal tree nodes were reconstructed by TreeTime (Sagulenko et al., 2018). We extracted the part of the alignment that corresponded to the S gene, and collapsed the internal tree branches without mutations in the S gene. The final tree had 1,783 internal branches. For each internal branch, we listed the amino acid mutations that occurred at this branch.

### Concordantly evolving site pairs

To study the concordant and discordant evolution of pairs of sites in SARS-CoV-2 spike, we applied our approach to the distribution of mutations in the S gene on the reconstructed SARS-CoV-2 phylogeny. 185 of the sites carried two or more mutations on internal tree branches. We considered all 17,020 unordered pairs of these sites.

We detected 45 concordantly evolving site pairs which comprised 46 sites (table S7, figure 1A, figure S3). Our phylogenetic approach for detecting concordantly and discordantly evolved site pairs relied on the assumption that the tree provided for analysis is correct. To check the robustness of our results to uncertainty of phylogenetic reconstruction, we repeated the analysis on the tree reconstructed for the same set of sequences by the USHER (Turakhia et al., 2021) method utilizing maximum parsimony approach (see Methods). Among the 45 site pairs inferred to be concordantly evolving (table S7), 33 were also concordantly evolving on the USHER tree at 50% FDR, including 28 at 10% FDR (table 1, figure 1A, table S8). Thus, we conclude that for 73% (33/45) of the detected concordantly evolved site pairs, the statistical signal was strong enough to be insensitive to phylogenetic uncertainty. We focus on the IQ-TREE results below.

**Figure 1.**
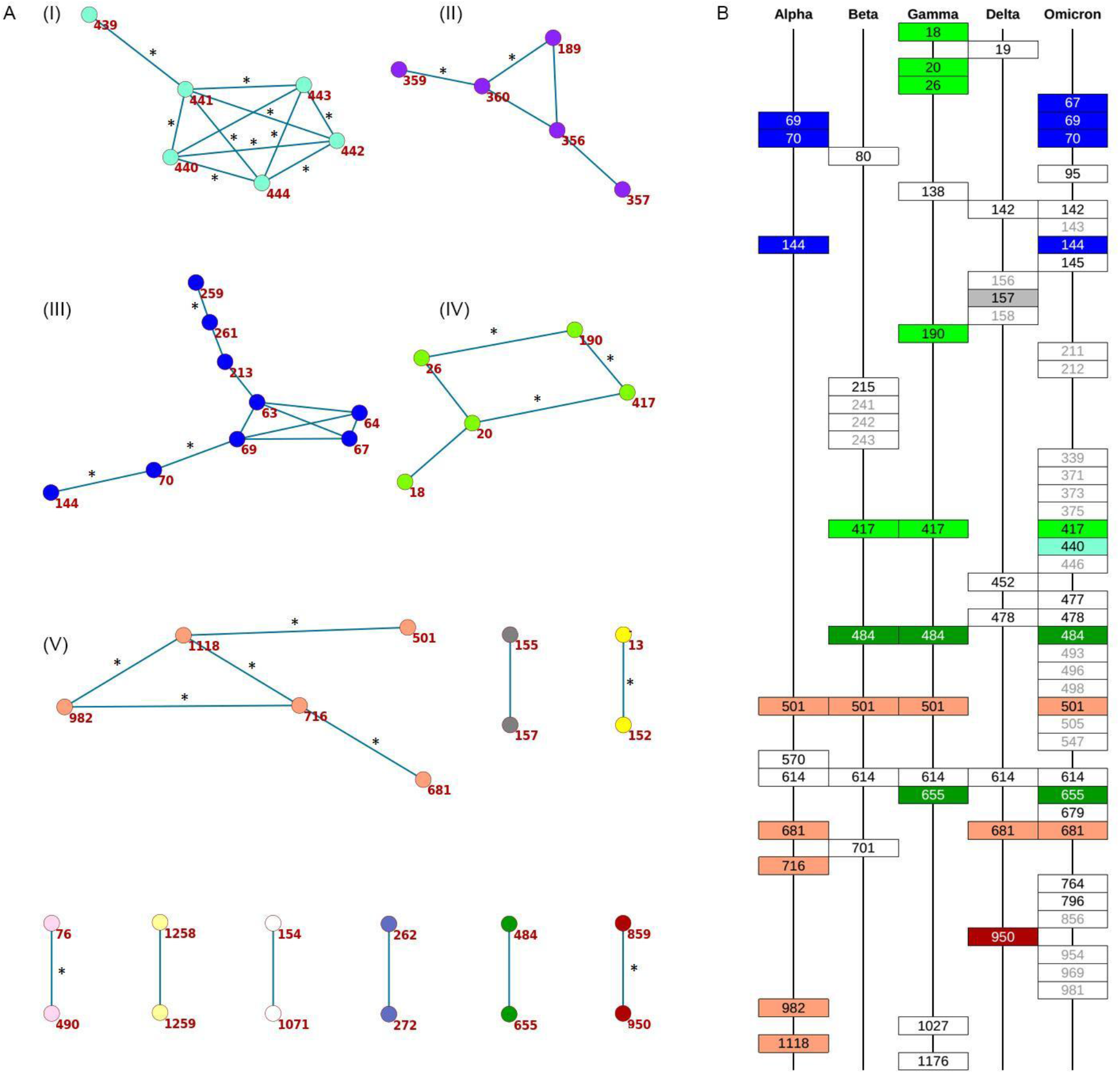
A. Clusters of concordantly evolving sites in spike protein. Graph vertices represent sites, and edges represent concordantly evolving pairs (table S7). The graph consists of 13 connected components, 8 of which contain just a single edge. Site pairs that were among the set of the best scoring pairs predicted for the alternative USHER (Turakhia et al., 2021) topology (FDR<10%, table 1) are marked by asterisks. Latin numbers are shown for referring to the site groups in the text. B. Concordantly evolving sites among the lineage-defining sites of Alpha, Beta, Gamma, Delta (B.1.617.2+AY.*) and Omicron (BA.1) VOCs (Emma B. Hodcroft., 2021). Concordantly evolving sites are colored in accordance with the clusters in panel A. Sites with fewer than 2 mutations which were not included in the analysis are in gray.

To characterize the detected concordantly evolving site pairs, we considered the positions of their sites on the S-protein trimer structure (PDB ID: 7JJJ). The mean distance between coevolving site pairs was shorter (20.17A) than expected for random site pairs between the 185 sites with two or more mutations on internal branches (36.71A, P=0.0004) as well as for random site pairs between the 46 sites involved in concordant evolution (36.64A, P<1E-4). Among the 42 coevolving site pairs for which the distances on the protein structure were known, in 18 (43%), the two sites were in contact, i.e., within 5A from each other.

To visualize the detected concordantly evolving site pairs, we plotted them as a graph where vertices represented the 46 sites, and edges represented the 45 pairs formed by them (figure 1A). The graph has 13 connected components; five of them are subgraphs each containing between 5 and 9 sites, and the remaining 8 each consisting of a single site pair. Sites of three of the five subgraphs with multiple vertices formed dense clusters on the protein structure, and for each such cluster, all or almost all sites belonged to the same domain; sites from the other two components were distributed dispersedly (figure 2A). Hereafter, we referred to these clusters of sites by latin numbers I-V. All six sites of the first dense cluster 439-444 (I) were in the receptor binding motif (RBM) within the receptor binding domain (RBD). The four sites of the second dense cluster (II) 356, 357, 359 and 360 were in RBD, while the fifth site 189 was in the NTD domain. Interestingly, in the open conformation (PDB ID: 7KL9), sites 357, 359 and 360 contacted the NTD domain of the adjacent subunit of the S-protein trimer, but in the closed conformation (PDB ID: 7JJJ) they were not in contacts (fig. 2A and 2B). The third dense cluster (III) 63, 64, 67, 69, 70, 144, 213, 259 and 261 was in the NTD domain in the region of binding of neutralizing antibodies (McCarthy et al., 2021). Four sites 18, 20, 26 and 190 in the first of the two dispersed clusters (IV) were in the NTD domain, and one site 417 was in the RDB. The second dispersed cluster (V) comprised sites from different domains: RBM – 501, near S1/S2 cleavage site – 681, S2 – 716 and 982 and in the connecting domain (CD) 7#x2013; 1118 (figure 3).

**Figure 2.**
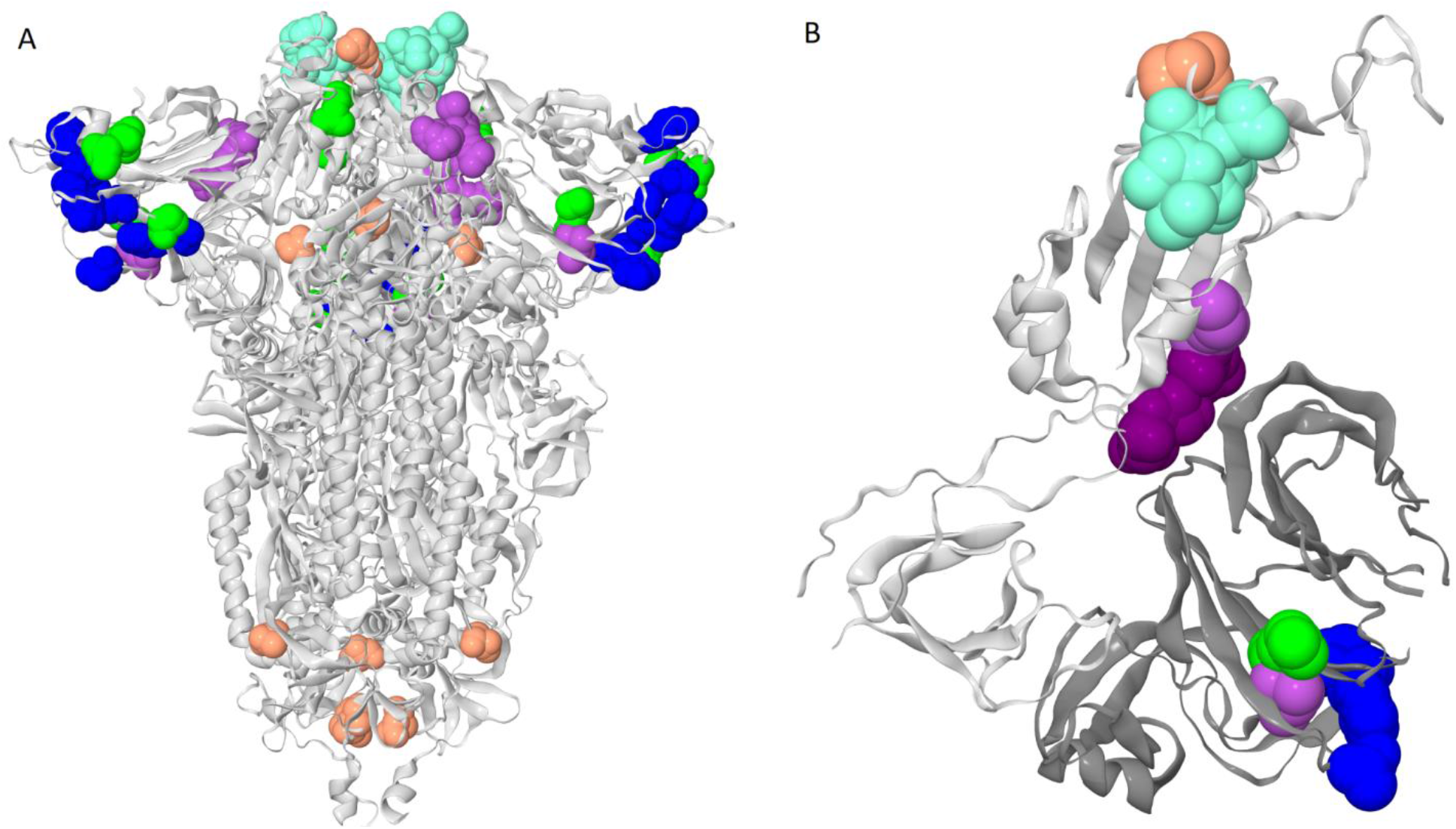
Clusters of coevolving sites on the protein structure. Sites of the five clusters that comprise multiple coevolving pairs of sites are shown as spheres, with color coding matching figure 1A. A. Closed conformation of the S-protein trimer (pdb: 7JJJ). B. Open conformation of the S-protein trimer (pdb: 7KL9): for clarity, only residues 320-590 of one subunit and NTD 14-303 of the adjacent subunit are shown. NTD is shown in dark gray; residues 357, 359 and 360 are shown in dark purple.

**Figure 3.**
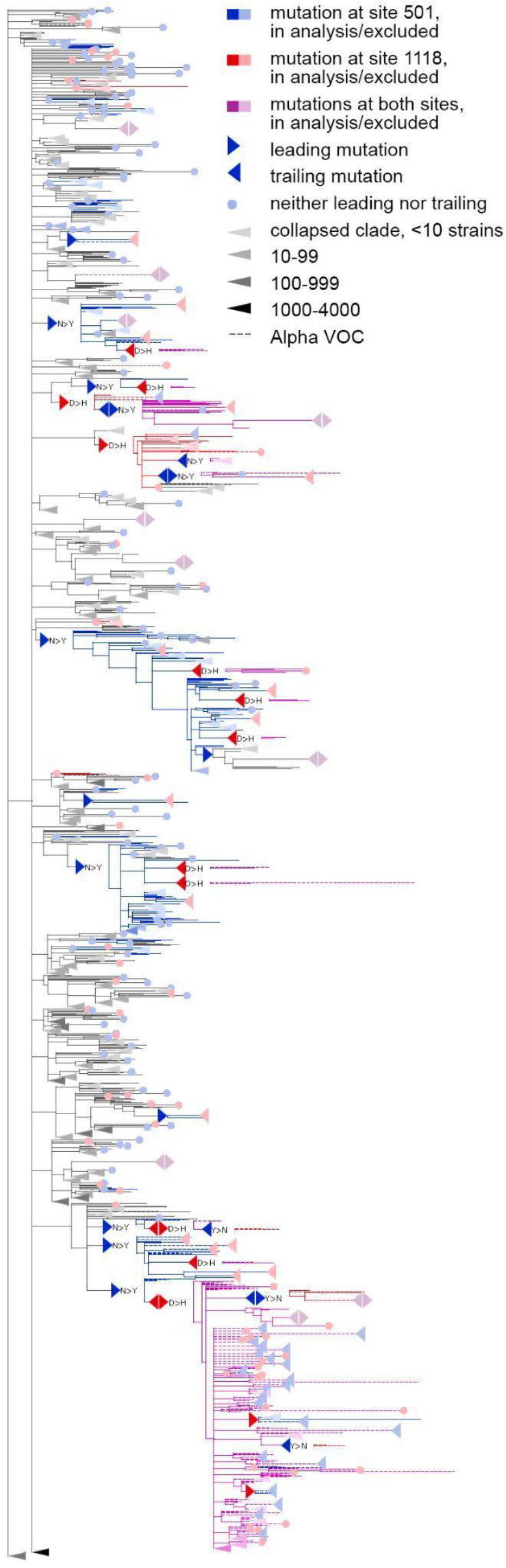
Concordantly evolving pair of sites 501 and 1118. The dashed branches correspond to the sequences of Alpha VOC. For clarity of presentation, some of the clades without mutations in these two sites are represented by long triangles, with intensity of color indicating the number of strains in the clade.

### Discordantly evolving site pairs

We detected four pairs composed of six sites that exhibited discordant patterns of evolution (table 2, figure S4). Three of these site pairs were among the 16 discordantly evolving pairs between the 15 sites detected for the alternative USHER topology for the same level of FDR (table S9). The signal of discordant evolution indicated that mutations at one site in a pair arose preferentially at a specific allelic context at another site; e.g. mutations Q675H and Q677H occurred mostly on the wild-type background N in the 501 (figure 4), while mutations A653V and S982A occurred mostly on the derived background G at site 614.

**Figure 4.**
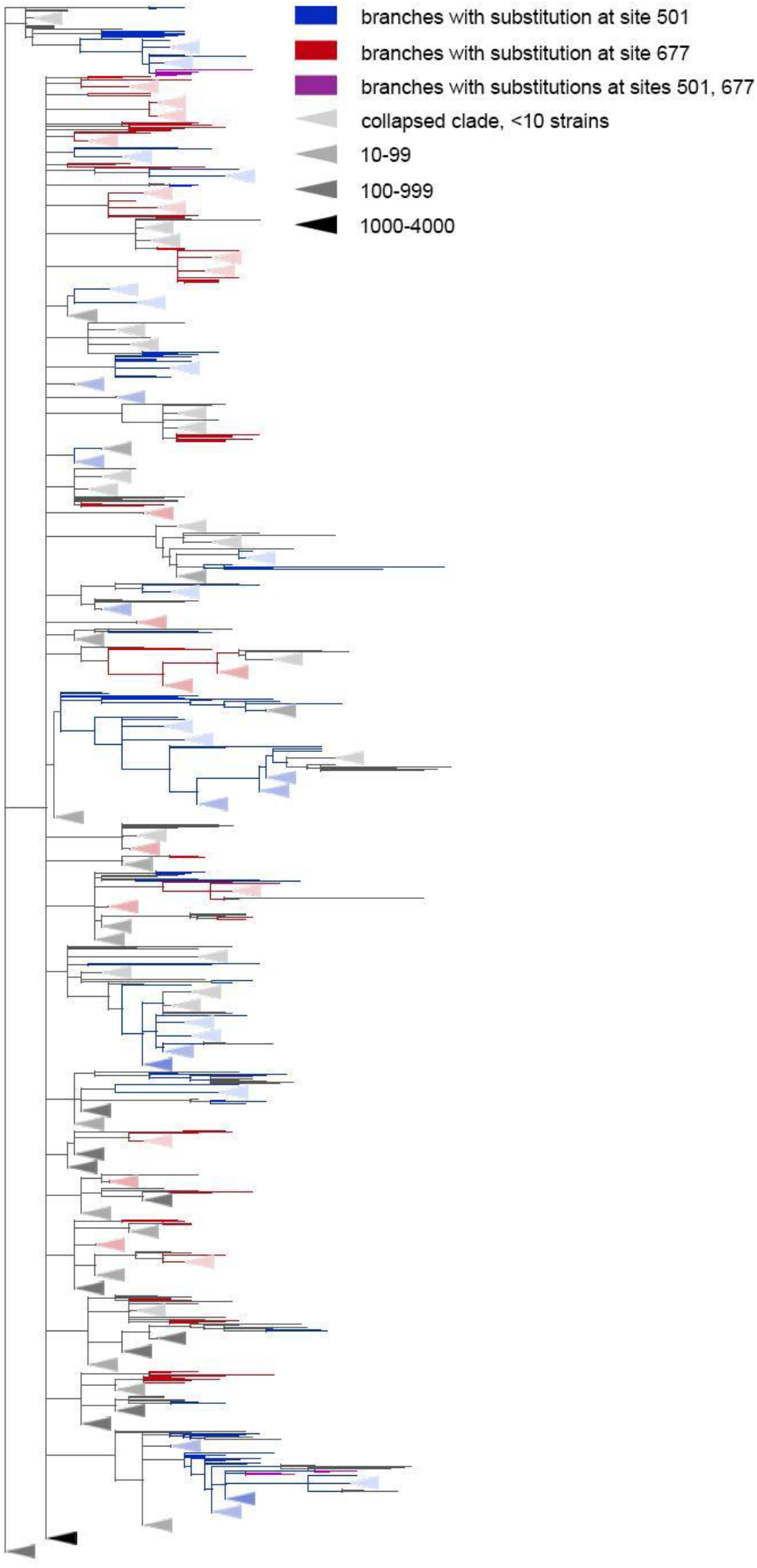
The Q677H avoids to occur on the background of N501Y. Branches carrying substitutions at site 501 are shown in blue; at site 677, in red; at both sites, in violet. Some clades without any analyzed mutations were truncated. Such clades are represented by long triangles, with intensity of color indicating the number of strains in the clade (light shade – less than 10 strains, medium shade – from 10 to 100, dark shade – from 100 to 1000, darkest shade – over 1000 strains).

**Table 2.**
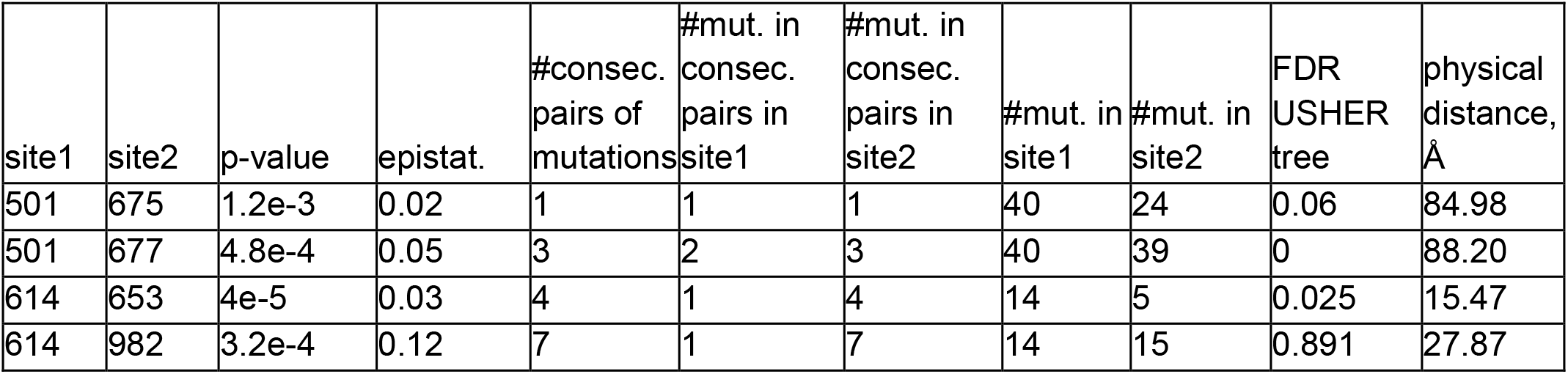
Discordantly evolving sites of the SARS-Cov-2 S-protein. The following characteristics are shown: coordinates on the S-protein sequence, nominal p-values, the value of the epistatic statistics, the total number of consecutive mutation pairs for the two corresponding ordered site pairs, numbers of mutations in consecutive pairs at site1 and site2, total numbers of mutations at site1 and site2, FDR value corresponding to the p-value of the site pair obtained for the alternative phylogeny reconstructed by USHER (Turakhia et al., 2021), and the distance in the protein structure (PDB ID: 7JJJ).

| site1 | site2 | p-value | epistat. | #consec. pairs of mutations | #mut. in consec. pairs in site1 | #mut. in consec. pairs in site2 | #mut. in site1 | #mut. in site2 | FDR USHER tree | physical distance, Å |
| --- | --- | --- | --- | --- | --- | --- | --- | --- | --- | --- |
| 501 | 675 | 1.2e-3 | 0.02 | 1 | 1 | 1 | 40 | 24 | 0.06 | 84.98 |
| 501 | 677 | 4.8e-4 | 0.05 | 3 | 2 | 3 | 40 | 39 | 0 | 88.20 |
| 614 | 653 | 4e-5 | 0.03 | 4 | 1 | 4 | 14 | 5 | 0.025 | 15.47 |
| 614 | 982 | 3.2e-4 | 0.12 | 7 | 1 | 7 | 14 | 15 | 0.891 | 27.87 |

### Coordinated evolution of sites carrying VOC mutations

Many of the concordantly evolving sites carried mutations that defined VOC lineages. In what follows, we discuss the sites of concordant mutations found among the sites carrying the characteristic mutations of specific VOCs (see figure 1B). We refer to sites bearing lineage-defining mutations as lineage defining sites.

For the Alpha VOC, eight (69, 70, 144, 501, 681, 716, 982 и 1118) out of the ten lineage defining sites (Emma B. Hodcroft., 2021) were among the 46 concordantly evolving sites. Three of these sites, 69, 70 и 144, were within the dense cluster III of sites located in the NTD; the remaining five sites represented the dispersed cluster V.

For the Beta VOC, three sites (417, 501 and 484) out of the ten lineage defining sites were among the concordantly evolving sites, but these sites did not form pairs with each other.

For the Gamma VOC, seven out of the twelve lineage-defining sites (18, 20, 26, 417, 484, 501 and 655) were among the concordantly evolving sites. Four of these sites (18, 20, 26 and 417) were within the dispersed cluster IV of sites. Two sites (484, 655) constituted a distinct cluster with a single pair (figure 5). Finally, site 501 had no concordantly evolving partners.

**Figure 5.**
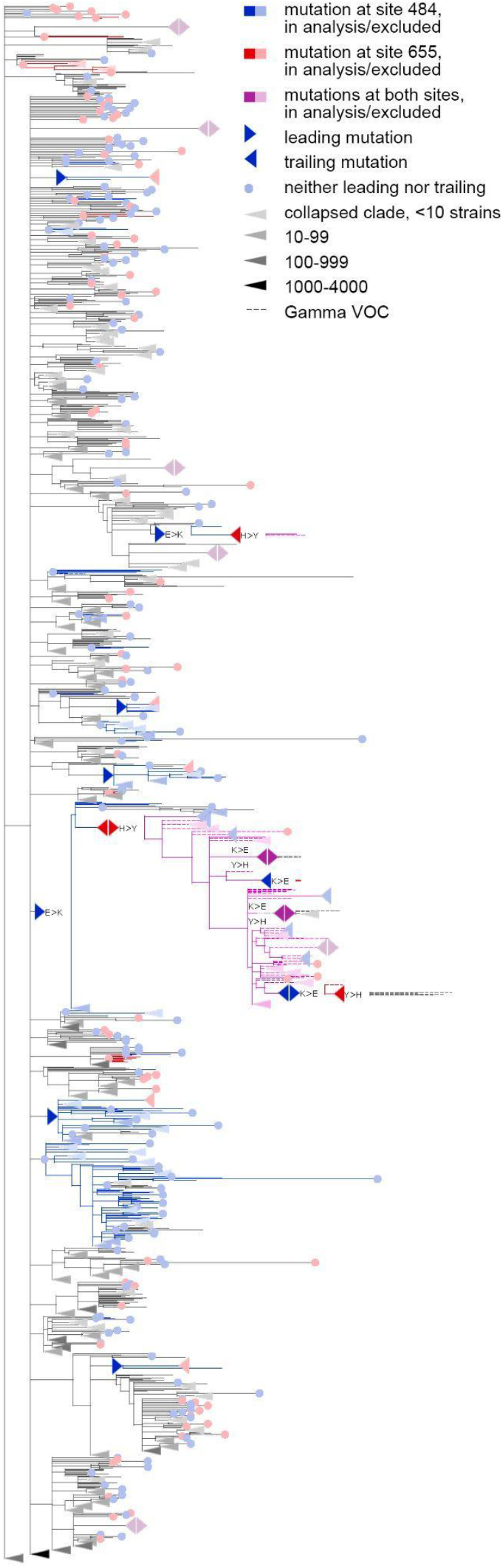
Coevolution of S-protein sites 484 and 655. The dashed branches correspond to the sequences of Gamma VOC. For clarity of presentation, some of the clades without mutations in these two sites are represented by long triangles, with intensity of color indicating the number of strains in the clade.

For the Delta VOC (B.1.617.2+AY.*), three out of the ten lineage defining sites (157, 681 and 950) were among the set of concordantly evolving sites, however, these sites belonged to different clusters and they did not form pairs with each other.

Finally, for the Omicron VOC (BA.1), ten out of the 36 lineage defining sites (67, 69, 70, 144, 417, 440, 484, 501, 655 and 681) were among the set of concordantly evolving sites. Notably, no Omicron sequences were in the dataset used to assess concordance, so the signal observed at these sites is not due to clustering of substitutions in them at the origin of Omicron. Seven of these ten sites, with the exception of the sites carrying deletions in Omicron (69, 70 and 144), previously were shown to evolve under positive selection (Martin et al., 2022). Among these ten sites, four (67, 69, 70 and 144) were within the dense cluster III of sites located in the NTD. The sites 501 and 681 belonged to the highly dispersed cluster V of sites but did not form a pair, suggesting that any interactions between them could be indirect and mediated by other sites; site 501 is located within the RBM, while site 681 is near the S1/S2 furin cleavage site (FCS). Two sites 440 and 417 each were the only representatives of the corresponding cluster and had no coevolving partners among the lineage-defining sites. Finally, the two remaining sites, 484 and 655, formed a pair of coevolving sites which represented a separate single-edge cluster (figure 1A and figure 5). Again, the first site in this pair was within the RBM and the second site was within the FCS. While 484A is the characteristic allele of the Omicron VOC, it was rare in previously circulating strains: indeed, mutation E484A occurred in just a few (8 out of 7348) of the terminal branches of our phylogenetic tree and always in the context of the ancestral histidine at site 655. As our analysis disregards substitutions at terminal branches, E484A thus could not have contributed to our epistatic statistic. Instead, the signal of epistasis between sites 484 and 655 was formed by other mutations, notably, the frequently occurring E484K.

These findings suggest that the lineage-defining sites are enriched in concordantly evolving site pairs. To formally test this, we generated 400 artificial datasets by randomly redistributing the mutations on the phylogeny while preserving the numbers of substitutions at each site and on each branch of the tree, and applied our method for inference of coordinated evolution for each such dataset. For each pair of sites in the real or in an artificial dataset, we had estimated the upper p-value as the probability to obtain the value of the epistatic statistic for independently evolving sites at least as high as that observed for the dataset. We ordered all site pairs according to their upper p-values in ascending order, and compared the difference of mean ranks of pairs of lineage-defining sites and mean ranks of other pairs of sites, both for the real and for the 400 random distributions of substitutions on the phylogeny (see Methods). All VOCs except Omicron had a stronger signal of coordinated evolution (lower ranks) for pairs of lineage-defining sites than for the remaining site pairs. This could not be explained by a difference in numbers of substitutions in lineage-defining and other sites because we have preserved numbers of substitutions at sites when generating datasets (tables S10-S14). Thus, our findings indicate that the lineage-defining sites of VOCs comprised pairs with a stronger signal of concordant evolution than expected.

As the lineage-defining sites of a VOC are by definition those that carry mutations at the origin of this VOC, the signal of concordant evolution at lineage-defining sites could arise from clustering of mutations at these sites at VOC origin. To address this, we asked whether concordant evolution of lineage-defining sites of a VOC is also observed when this VOC itself is excluded from analysis. For this, we separately pruned all isolates belonging to each of the four VOCs Alpha, Beta, Gamma and Delta from the tree. Note that our datasets did not contain Omicron from the start. For each of four pruned trees, we separately predicted the concordantly evolving site pairs, and tested for enrichment of pairs of lineage-defining sites. For Alpha, the pairs of lineage-defining sites still had a stronger signal of concordant evolution, indicating that at least for this VOC, the observed concordance is not due to clustering of substitutions at VOC origin (table S15). For the other three VOCs, Beta, Delta and Gamma, no enrichment was detected (tables S16-S18). In fact, for Gamma, after pruning of the VOC clade, pairs of lineage-defining sites became depleted among the pairs with a stronger signal of concordant evolution (P=0.0272, table S18); in this VOC, the observed signal of concordant evolution of lineage-defining sites is mainly caused by reversions of lineage-defining mutations within the VOC clade (figure 5).

## Discussion

In this work, we modified the phylogenetic approach for detection of interdependently evolving pairs of sites that had been previously successfully applied for influenza A (Kryazhimskiy et al., 2011; Neverov et al., 2015) and mitochondrial proteins (Neverov et al., 2021), and applied it to the Spike protein of SARS-CoV-2. Using simulations, we show that our revised method has a better specificity and sensitivity for detection of epistatically interacting site pairs than its original version, and that it is able to detect positive as well as negative epistasis between sites. Our simulations involved multiple sites with unfavorable alleles, so there were multiple adaptive mutations, allowing competition between multiple clones and hitchhiking. Despite these confounders, our method did not produce spurious signal of epistasis when epistasis was not a part of the simulation (tables S1 and S2, figures S1 and S2) and was able to detect true interacting site pairs with reasonable FDR for the epistatic mode of evolution of genotypes (tables S3 and S4).

Similar to other methods for inference of factors of evolution from comparative genomics data, the accuracy of our method depends on the validity of its assumptions. First, we assume that the observed changes in the ancestral reconstructed states between adjacent nodes of the phylogeny correspond to actual mutations rather than being artefactual. Unfortunately, for large-scale sequencing projects such as that of SARS-CoV-2, some extent of mistakes in the called sequences is unavoidable, and these mistakes are not random. Specifically, as was noted previously (Martin et al., 2022), the propensity to call the ancestral nucleotides (reference bias), particularly at sites of low NGS read coverage, may lead to reversions to wild-type alleles, in particular for lineage-defining mutations. Indeed, the main cause of calling of wild-type alleles at multiple sites is the selective preference of PCR or sequencing primers to some genotypes that leads to a high variation in coverage along the genome of different isolates. In periods of change of the dominant variant, two genetically different variants circulate with high population frequencies, and mixed infections or cross-contaminations of samples may result in artifactual hybrid sequences. This can be a problem for our method. Exclusion of terminal tree branches mitigates these problems by focusing the analysis on more frequent genotypes because spurious reversions more likely occur in the tree on terminals. This problem is also partially addressed by the fact that we count all trailing mutations following a single leading mutation as one, so our method encourages site pairs with multiple phylogenetically unrelated pairs of consecutive mutations; this requirement decreases the impact of a systematic loss of mutations at some sites of specific variants of genotypes.

Second, our approach relies on the correctness of the phylogeny. The inference of the true phylogeny for SARS-Cov-2 is difficult due to the huge amount of data and limited sequence diversity (Morel et al., 2021; Rochman et al., 2021b). To assess the impact of phylogenetic uncertainty on the results of analysis, we compared the sets of concordantly and discordantly evolving site pairs inferred for two trees: the maximum likelihood (ML) tree obtained by IQ-TREE 2 (Minh et al., 2020) and maximum parsimony (MP) tree obtained by USHER (Turakhia et al., 2021). The results were relatively stable (tables 1 and S7, S8), indicating that most of the detected signal is robust to the method of phylogenetic reconstruction.

Third, we assume that evolution is clonal, so that the phylogenetic tree reflects the true evolutionary history of the genome. If genotypes recombine, some genomic sites would be incompatible, i.e., have different evolutionary histories (Bruen et al., 2006). On the whole genome phylogenetic tree, those sites with evolutionary history disagreeing with the majority would be enriched in spurious parallel or convergent substitutions, possibly affecting the signal of concordance. While the recombination frequency for SARS-CoV-2 is unknown, it has been estimated that about 3-5% of circulating genotypes are recombinants (Kozlakidis, 2022; Turkahia et al., 2021) and the frequency of occurrence of recombination breakpoints in the S gene is up to three times higher than in the rest of the genome (Turkahia et al., 2021). While our dataset included four sequences of the XB recombinant lineage, exclusion of these sequences did not affect our results. Nevertheless, some of the recombinants could remain unannotated, particularly those including just a few samples and/or originating from similar sequences.

The signals of concordant and discordant evolution that we observe could come from at least two natural sources. Firstly, they could arise from epistatic interactions between sites, an explanation which we favored in our previous works (Kryazhimskiy et al., 2011; Neverov et al., 2021, 2015). Previously, Ruan et al. (Ruan et al., 2022) interpreted the stepwise accumulation of mutations at origin of Gamma and Delta VOCs as evidence for epistatic interactions between lineage defining mutations. Under this explanation, concordant evolution of a pair of sites is indicative of positive epistasis, and discordant evolution, of negative epistasis. Recently, epistasis in the Spike protein has been demonstrated experimentally: at 15 of the RBD sites, the effects of mutations were shown to change on the background of N501Y (Starr et al., 2022). Unfortunately, it is hard to cross-validate the epistatic interactions between the evolutionary and experimental analyses. In our dataset, these sites are conserved: for 14 out of the 15 sites, there are no mutations on internal tree branches, making them unfit for our analysis. The remaining site 449 carried a single mutation Y449H which by itself is known to strongly decrease ACE2 binding affinity (Starr et al., 2022); this mutation co-occurred with N501Y on one internal branch which had two descendent leaves, leading to a p-value of 0.18 for this pair in our test.

Our observation that lineage-defining sites of Alpha evolve concordantly even outside of the Alpha clade supports the significance of epistasis at origin of at least this VOC (table S15). Another piece of evidence comes from the observation of discordant evolution at sites carrying mutations which are likely individually beneficial. Among the four predicted discordantly evolving pairs, two pairs (501, 677) and (501, 675) are between the three sites whose effects of mutations were assessed experimentally or computationally (figure 4). The N501Y mutation increases the binding affinity of Spike to ACE2 up to 15-fold (Starr et al., 2022) and increases infectivity (Liu et al., 2022); the Q677H mutation increases infectivity, propensity to syncytium formation and escape of neutralization by serum of vaccinated people (Zeng et al., 2021, p. 2); and the Q675H mutation is predicted to increase furin binding affinity (Bertelli et al., 2021). Although experiments suggested a positive effect of Q677H on the background of VOCs Alpha and Gamma both carrying N501Y (Zeng et al., 2021, p. 2), this fact is in disagreement with very low population frequencies of Q677H in these VOCs (Gangavarapu et al., 2022a, 2022b, 2022c). The same is true for the Q675H mutation: while it was observed in some isolates of VOCs with the N501Y lineage-defining mutation, the population frequencies of these strains were also very low (Bertelli et al., 2021).

The second natural source of coordinated occurrence of mutations is differences in selection regimes between tree branches. Over the course of the pandemic, selection on the virus has changed due to the dynamics of herd immunity of the human population, with selection favoring immune escape mutations increasing with time. In particular, this was proposed to underlie the evolution of Gamma (Gräf et al., 2021, p. 1) and Omicron (Martin et al., 2022) VOCs. Additionally, distinct selection regimes could correspond to long-term infection in immunocompromised individuals or in non-human hosts under reverse zoonosis events. Overall, we find that the VOCs at their origin have gained mutations at concordantly evolving sites (tables S9-S12). Interestingly, the set of the VOCs that we indicate as evolving in a coordinated manner, Alpha, Gamma and Omicron, are the same three VOCs that are characterized by an increase in the rate of evolution at their origin and therefore have been suggested to had evolved in distinct evolutionary regimes, perhaps that of an immunocompromised patient (Hill et al., 2022; Martin et al., 2022). A possible explanation of constellations of lineage-defining mutation at the origin of VOCs is that selection in favor of these mutations increases in immunocompromised patients, leading to “episodes” of adaptation (Martin et al., 2022) during which the rate of their accumulation is increased compared to the baseline observed in the general population, making them clustered in the branches corresponding to such patients. This mechanism would lead to a signal of concordance in our analysis (Neverov et al., 2021). However, it is unclear why the virus had to “wait” for an immunocompromised individual to evolve the mutations that also increased its fitness in the general population. Also, we do not expect such episodic selection to lead to signals of discordant evolution.

Finally, the signal of concordant evolution at the origin of VOCs could come from the combination of both factors: a distinct selection regime and epistasis. The role of immunocompromised patients in the evolution of SARS-Cov-2 is even higher for epistatic than for the non-epistatic mode of evolution (Smith and Ashby, 2022). If the viral fitness, as it has been previously proposed (Hill et al., 2022; Rochman et al., 2021a; Smith and Ashby, 2022), is a trade-off between transmission efficiency and ability to avoid herd immunity, and the transmission component of the fitness landscape has valleys of low-fitness genotypes, the immunocompromised individuals with prolonged infectious periods due to relaxed selection for transmission efficiency allow the virus to accumulate mutations and cross these valleys. Direct experimental studies of the possible epistatic interactions between coevolving sites will help elucidate the mechanism of origin of radically novel viral variants.

## Materials and Methods

### Constructing the set of sequences

Masked full genome sequence alignment and corresponding metadata were downloaded from gisaid.org on 07.09.2021 comprising sequences for 3,299,439 isolates table S19). Based on metadata, all sequences from nonhuman hosts, without collection dates, or with wrongly formatted collection dates were excluded. For each sample, the number of gaps, ‘N’s and ambiguous characters were computed for each genome sequence and separately for the S-gene in non-masked positions. For each sequence, we calculated the number of positions that contained a nongap, non-‘N’ symbol that were aligned to non-gap positions of the reference genome sequence WIV04. We excluded sequences from the analysis with fewer 29,000 aligned positions, or with <95% of sequence length in aligned positions. Next, we sorted sequences by sampling dates and then converted each sequence into a list of changes relative to the reference, treating consecutive gaps as one change. Then, we excluded sequences having too many changes for their sampling dates. For that, for each sampling date, we computed the mean and standard deviations of the number of changes for all sequences whose sampling dates were within a half year time interval centered at this date. All samples with the number of changes exceeding the mean value by two standard deviations or more in the corresponding time interval centered on the sampling date were filtered out. Samples with preliminary stop codons inside the S gene were also excluded. This left us with 2,676,884 sequences for further analysis.

We partitioned the remaining sequences into groups of equivalence such that all sequences in each group had the same list of mutations in the S gene, and selected the sequence with the earliest collection date as the class representative. To further cluster the sequences, we reimplemented the UCLUST (Edgar, 2010) algorithm in a custom python script to allow it to process the huge amount of SARS-CoV-2 data. In contrast to the original UCLUST implementation that receives nucleotide or amino acid sequences as input, our script clustered the lists of changes in sequences that occurred relative to the sequence of the reference genome. This sped up computation, because there were few changes in SARS-CoV-2 sequences compared to the genome length. The pairwise distance between the two lists of changes was defined as the number of changes unique to each list. At the start of the procedure, samples were ordered by the sampling date; the first sample was defined to be the centroid of the first cluster and added to the list of centroids. Next, the remaining samples were iteratively compared with the centroids in the list: if the distance to some centroid was less than three, the sample was added to the corresponding cluster, otherwise it was added as a new entry to the list of centroids. Thus, by construction, cluster centroids tend to be the earliest representatives of their members. For each cluster, all samples from the corresponding groups of equivalent sequences were pooled together, and the sample having the highest quality sequence was selected as the representative of the cluster. The highest-quality sequence was defined as the sequence having the minimal number of gaps or ‘N’ characters in the S gene. If several sequences complied with the previous condition, the sequence with the minimal number of ambiguous characters in the S gene was selected as the representative of the cluster. If there were more than one such sequence, the one additionally having the minimal number of gaps or ‘N’ characters in the whole genome sequence was selected. If still more than one sequence met all the previous conditions, the first of them which had the minimal number of ambiguous characters in the whole genome sequence was finally chosen. All gaps in the selected complete genome sequences were converted to reference characters, and insertions relative to reference sequence were ignored.

### Phylogenetic analysis and inference of ancestral sequences

The phylogenetic tree was constructed with IQ-TREE (v. 2.1.2, model = GTR+I+G) (Minh et al., 2020). Additionally, we obtained an alternative topology for these sequences by USHER (Turakhia et al., 2021); for this, we inserted the sequences into a prebuilt global SARS-Cov-2 phylogeny of publicly available genome sequences (http://hgdownload.soe.ucsc.edu/goldenPath/wuhCor1/UShER_SARS-CoV-2/public-latest.all.masked.pb.gz accessed on 10.01.2022). Some of our selected GISAID sequences had already been in the global tree, so we inserted only those which were absent there. Finally, we extracted the subtree corresponding to the analyzed sequences from the global tree.

The trees were rooted by the outgroup sequence USA-WA1/2020 (EPI_ISL_404895). For both trees, ancestral sequences were reconstructed by TreeTime (v. 0.8.2) (Sagulenko et al., 2018) with default parameters. Because we focused on the evolution of the S gene, we removed from each tree the internal nodes which had S-gene sequences identical to their parental nodes. Finally, we obtained the list of mutations for each tree branch.

### Concordantly and discordantly evolving pairs of sites

Our approach to detection of concordantly and discordantly evolving pairs of sites is a development of the phylogenetic method published earlier (Kryazhimskiy et al., 2011; Neverov et al., 2021, 2015). It is based on counting consecutive pairs of mutations on the branches of a phylogenetic tree. A pair of mutations at two different sites is called consecutive if a mutation in one of the sites occurs in the subtree of the branch which carries a mutation in the other site, and there are no other mutations at these sites on the branches that constitute the path between them (Kryazhimskiy et al., 2011). Here, we consider four models for detection of epistasis which are based on this approach. Two of these models account for identities of ancestral and derived amino acids changes by the mutations at sites; the other two models disregard the identities of the mutations. We define the epistatic statistics in a general form which is used for all four models. The expression of the epistatic statistic for models that ignore identities of alleles could be straightforwardly obtained from the general form.

For an ordered pair of sites *(i,j),* the epistatic statistics *e_(i,j)_,* is the weighted number of consecutive pairs. It is formally defined as

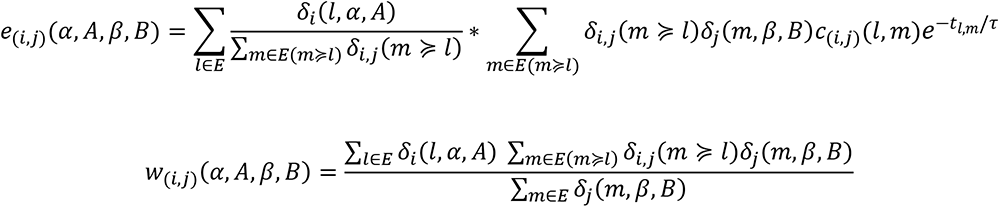

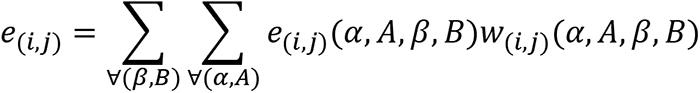

Here, *α* and *β* are the ancestral alleles, and and are the derived alleles at sites *i* and *j*; *E* is the set of all tree edges; and 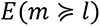 is the set of edges descendant to the edge *l*. Indicator functions 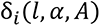 and 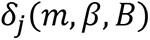 equal to one if on branches *l* and *m,* mutations at sites *i* and *j* occur from the specified ancestral alleles to the specified derived alleles. The indicator function 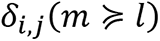 equals to one if a mutation occurs at branch *l* at site *i,* and a consecutive mutation occurs at branch *m* at site *j*. is the length of the shortest path between tree branches *l* and *m*. The function *c_(i,j)_(l,m)* accounts for possible incomplete time resolution of the sequence of occurrence of mutations: it equals one if mutations at sites *i* and *j* occur on different branches, 0.5 if mutations at both sites occur on the same branch *l=m*, 1.5 if mutations at both sites occur at the same branch *l* or *m* and are followed or preceded by a mutation at one of the sites *i* or *j* at another branch, and 0.25 if mutations at both sites occur on both branches *l* and *m*. The weight *w_(i,j)_(α, A, β, B)* is the fraction of all mutations from *β* to at site *j* that occur on the background of the mutation from *α* to at site *i*. In the models of detection of epistasis which ignore allele identities, the weights are set equal to 1. The epistatic statistic for an unordered pair is a total of two statistics for ordered pairs: *e_i,j_* = *e_(i,j)_* + *e_(j,i)_*.

We used two different null models for the epistatic statistics. The first model randomly and independently reshuffles mutations at each site; the second model specifically accounts for distribution of mutations at each ordered pair of sites (*i,j*). To do that, the phylogenetic positions of mutations at the first site of a pair (background site) *i* are considered fixed, and the positions of mutations at the second (foreground) site *j* are reshuffled, thus unlinking the background and foreground. For both null models, reshuffling of mutation positions between tree branches preserves the total number of mutations at each site and the total number of mutations at each branch by using BiRewire utility (Gobbi et al., 2014). Combinations of two variants of epistatic statistics (with and without alleles) and two variants of the null model (with linked and with unlinked background and foreground) provide four models for detection of site pair epistasis which are compared in this study.

For the two models that account for allele identities, we generate amino acid sequences for internal and terminal nodes of the tree. For that, starting from the root sequence and traversing the tree from root to tips, we generate derived alleles for each mutation on a tree branch from the empirical allele distributions at this site, conditioned on the allele at the parental node of the branch. For the null model with unlinked background and foreground, the sequences in the tree nodes for the background remain unchanged, while for the foreground, new sequences are generated. For generating random distributions of mutations on tree branches and for calculation of epistatic statistics, we used the Bio::Phylo Perl module (Vos et al., 2011) for traversing phylogenetic trees.

For each model, we perform 50,000 permutations of positions of mutations; for the two allele-aware models, for each permutation, we also generate the amino acid sequences at tree nodes. For each permutation, we calculate the epistatic statistics for unordered pairs, together with two p-values: the fraction of statistics equal or greater than the observed value (upper p-value), and the fraction of statistics equal or less than the observed value (lower p-value).

To account for multiple testing, we estimated the false discovery rates (FDR). For this, we randomly selected 400 out of 50,000 permutations. For each of the 400 permutations, for each unordered pair of sites, we calculated the epistatic statistic and the upper and the lower p-values. For each p-value threshold, we calculate the corresponding number of findings for the real dataset (R – declared positives) and the average number of findings in the fake dataset (E[V] – false positives). The FDR is the ratio of E[V] to R.

### Simulation of independent evolution of sites

To model independent evolution of sites, we used genome-wide forward simulator MimicrEE2 (Vlachos and Kofler, 2018). Under independent mode of evolution, MimicrEE2 multiplies the fitness changes caused by individual mutations to get the fitness of a genome. The initial population consisted of 50,000 identical haploid genotypes with 100 biallelic (*a* or *A*) sites. At the start of simulation, 20 sites were under positive selection, since the initial allele was deleterious: its fitness was equal to 0.9945, while fitness of the other variant was equal to 1. Another 20 sites evolved under negative selection, since the initial allele was beneficial. The remaining 60 sites evolved neutrally, with the two possible variants at each site having fitness of 1. We simulated evolution of the population for 5,000 generations, with mutation rate 5e-4 mutations per site per generation. Each 250 generations, 50 genotypes were sampled from the population, resulting in 1,000 sequences. These were further used to reconstruct the phylogeny and measure the signal of epistasis. For each of the four detection models, the minimal FDR threshold was obtained, such that if the desired level of FDR would be below the threshold, no false concordantly evolving pairs of sites would be predicted.

### Simulation of positively and negatively epistatically evolved site pairs

MimicrEE2 allows modeling of epistatic interaction between a pair of sites, assigning fitness values to all possible combinations of binary variants at these sites (*aA*, *aA*, *Aa* and *AA*). The fitness of a genome in this case is the product of fitness values of individual changes and fitnesses of variant combinations for specified pairs. The initial population consisted of identical genotypes with lowercase alleles at each site, with a total of 100 sites: 20 sites (or 10 pairs of sites) in positive epistasis, 20 sites in negative epistasis and 60 neutrally evolving sites with no epistatic interactions. At neutrally evolving sites, all variants had fitness equal to 1. To model positive epistasis between a pair of sites, we assigned fitness 1 to variant combinations *aa* and *AA* and fitness 0.9945 to *aA* and *Aa*, so that the first mutation at one of the sites was deleterious, and the consequent mutation at the second site restored the initial fitness. To model negative epistasis between a pair of sites, we assigned fitness 1 to variant combinations ab, aB and Ab, and fitness 0.8 to AB, so that the first mutation at one of the sites was neutral, and the consequent mutation at the second site was deleterious. Again, we simulated evolution of a haploid population of size 50,000 for 5,000 generations, with mutation rate 5e-4 mutations per site per generation, sampled 50 genotypes each 250 generations, and used the resulting 1,000 sequences to reconstruct the phylogeny and measure the signal of epistasis.

### Comparing the signal strength of coordinated evolution across site subsets

To compare the strength of concordant evolution for pairs of sites belonging to a specified site subset with other pairs of sites, we designed a test comparing the average ranks of pairs of sites in these two subsets of pairs. First, we ordered all site pairs from high to low strength of concordant evolution of their sites according to ascending upper p-values. Then, we calculated the mean ranks of site pairs in each of the two subsets: in a subset of pairs for which both sites belong to the specified subset of sites, and in the complementary subset of pairs. A direct comparison of ranks in these two subsets of pairs may be misleading, because a test for coordinated evolution of sites may assign better p-values to pairs of sites having some properties, e.g. those with higher evolutionary rates, which could be overrepresented in a specified subset of sites. Therefore, we need to compare the mean ranks for the bipartition of site pairs for the actual data with the distributions of mean ranks for the same bipartition for data simulating independent evolution of sites. As simulated data, we used a set of 400 permutations of mutations on the tree used for the FDR estimation. Calculating the ranks of pairs, we then assigned the average values of ranks for site pairs having the same values of the ordering statistic. As the test statistic, we used the difference of the mean ranks of pairs in two parts of the bipartition. To calculate the p-value of the test, the test statistic for the actual data was compared with the test statistic obtained for the simulated data.

We applied this procedure to separately tested subsets of lineage-defining sites of VOCs that circulated before May 2021: Alpha (B.1.1.7+Q.*), Beta (B.1.351.*), Delta (B.1.617.2+AY.*) and Gamma (P.1.*). The list of lineage-defining mutations in the S-gene was compiled according to (Emma B. Hodcroft., 2021) (accessed 23.07.2022). We considered only missense lineage-defining mutations and excluded from the analysis the site S:614, because the mutation D614G was fixed in all considered VOCs.

To test whether the enrichment of pairs of lineage-defining sites for Alpha, Beta, Delta and Gamma among concordantly evolving site pairs is due to mutations that occurred at origins of these VOCs and/or reversions of these mutations within the VOC clades, for each VOC, we obtained a pruned tree. For this, we removed from the phylogeny the isolates that were assigned by PANGOLIN (O’Toole et al., 2021) to the corresponding VOC in their metadata as well as the isolates of other lineages descendant to the ancestral node of the VOC that carried on its branch the most recent lineage-defining mutation. For each pruned tree, we then applied the procedure of finding concordantly evolving pairs of sites, and then the procedure of comparing the strength of concordant evolution for pairs of lineage-defining sites and the complementary subset of pairs.

### Data Availability

The codes and data files required to reproduce the analysis are available at https://github.com/gFedonin/EpiStat.

The data for reproducing analyses from the paper are in the archive – “sars-2.epistat.data.tgz”.

## Supplementary materials

**Table S1.** . Levels of spurious signal of concordant evolution, inferred in the simulated dataset with no epistasis by four variants of the method. Number of pairs, detected under p-value threshold 10^-4^, and estimated FDR for this threshold are shown.

| Model |  | #FP | min p-value | Maximal FDR threshold for #FP = 0 |
| --- | --- | --- | --- | --- |
| consider alleles | shuffle mutations in fgr. only | 2 | <1e-4 | 0,25 |
|  | shuffle mutations both in bgr. and fgr. | 3 | <1e-4 | 0,15 |
| ignore alleles | shuffle mutations in fgr. only | 4 | <1e-4 | 0,11 |
|  | shuffle mutations both in bgr. and fgr. | 24 | <1e-4 | 0,02 |

**Table S2.** Levels of spurious signal of discordant evolution, inferred in the simulated dataset with no epistasis by four variants of the method. Number of pairs, detected under p-value threshold 10^-4^, and maximal threshold on estimated FDR that allows to avoid false discoveries are shown.

| Model |  | #FP | min p-value | Maximal FDR threshold for #FP = 0 |
| --- | --- | --- | --- | --- |
| consider alleles | shuffle mutations in fgr. only | 2 | <1e-4 | 0,2 |
|  | shuffle mutations both in bgr. and fgr. | 19 | <1e-4 | 0,026 |
| ignore alleles | shuffle mutations in fgr. only | 4 | <1e-4 | 0,125 |
|  | shuffle mutations both in bgr. and fgr. | 16 | <1e-4 | 0,031 |

**Table S3.** . Numbers of truly and falsely predicted concordantly evolving pairs of sites for the simulated data with positively and negatively epistatically interacting sites. The evolution of the population of genotypes with twenty independently evolving pairs of epistatically interacting sites was simulated by MimicrEE2. Ten pairs of sites were evolving under recurrent positive epistasis and other ten pairs were evolving under magnitude negative epistasis. The four different detection models were applied and for each model the number of true (#TP) and false (#FP) predictions are shown for estimated FDR≤10%.

| Model |  | #FP | #TP |
| --- | --- | --- | --- |
| consider alleles | shuffle mutations in fgr. only | 5 | 10 |
|  | shuffle mutations both in bgr. and fgr. | 55 | 6 |
| ignore alleles | shuffle mutations in fgr. only | 22 | 8 |
|  | shuffle mutations both in bgr. and fgr. | 250 | 7 |

**Table S4.** Predicted concordantly evolving pairs of sites for the simulated data with positively and negatively epistatically interacting sites. The characteristics of predicted site pairs for the estimated FDR≤10% are shown: positions of sites on the primary sequence of S-protein (site1 and site2), the upper p-values (p-value), values of the epistatic statistic (epistat), numbers of consecutive pairs of mutations (#consec. pairs of mutations), numbers of mutations in consecutive pairs of sites (#mut. in consec. pairs in site1 and site2), total numbers of mutations on the internal tree branches in sites (#mut. in site1 and site2), the expected value of epistatic statistics (exp. epistat) and the corresponding standard error (SE). The true predictions are highlighted in bold.

| site 1 | site 2 | p-value | epistat | #consec. pairs of mut. | #mut. in consec. pairs in site1 | #mut. in consec. pairs in site2 | #mut. in site1 | #mut. in site2 | exp. epistat | SE |
| --- | --- | --- | --- | --- | --- | --- | --- | --- | --- | --- |
| **1** | **2** | **<1e-4** | **12,36** | **66,5** | **52** | **65** | **2** | **96** | **3,97** | **0,87** |
| **3** | **4** | **<1e-4** | **17,79** | **94** | **61** | **104** | **4** | **118** | **6,55** | **1,23** |
| **5** | **6** | **<1e-4** | **13,87** | **54** | **48** | **54** | **6** | **105** | **3,27** | **0,83** |
| 5 | 43 | <1e-4 | 21,43 | 108,5 | 82 | 100 | 43 | 105 | 12,6 | 1,89 |
| **7** | **8** | **<1e-4** | **11,36** | **50** | **42** | **50** | **8** | **109** | **3,22** | **0,78** |
| **9** | **10** | **<1e-4** | **17,09** | **90** | **64** | **88** | **10** | **117** | **6,62** | **1,19** |
| **11** | **12** | **<1e-4** | **13,46** | **53,5** | **48** | **51** | **12** | **108** | **2,54** | **0,69** |
| **13** | **14** | **<1e-4** | **11,63** | **52,5** | **43** | **56** | **14** | **106** | **2,85** | **0,72** |
| **15** | **16** | **<1e-4** | **11,84** | **69** | **47** | **70** | **16** | **116** | **5,88** | **1,08** |
| **19** | **20** | **<1e-4** | **9,06** | **47** | **39** | **47** | **18** | **109** | **4,29** | **0,88** |
| 48 | 68 | <1e-4 | 29,11 | 171,5 | 110 | 169 | 20 | 95 | 2,6 | 0,69 |
| **17** | **18** | **2e-4** | **8,23** | **66** | **38** | **65** | **31** | **95** | **6,61** | **1,23** |
| 19 | 31 | 2e-4 | 11,58 | 78,5 | 56 | 78 | 67 | 176 | 20,72 | 2,09 |
| 50 | 94 | 2e-4 | 24,34 | 162,5 | 100 | 151 | 68 | 183 | 17,55 | 1,9 |
| 78 | 94 | 2e-4 | 28,50 | 172 | 110 | 169 | 94 | 178 | 20,5 | 2,07 |

**Table S5.** . Numbers of truly and falsely predicted discordantly evolving pairs of sites for the simulated data with positively and negatively epistatically interacting sites. See legend for Table S3.

| Model |  | #FP | #TP |
| --- | --- | --- | --- |
| consider alleles | shuffle mutations in fgr. only | 4 | 7 |
|  | shuffle mutations both in bgr. and fgr. | 62 | 5 |
| ignore alleles | shuffle mutations in fgr. only | 4 | 9 |
|  | shuffle mutations both in bgr. and fgr. | 7 | 5 |

**Table S6.**
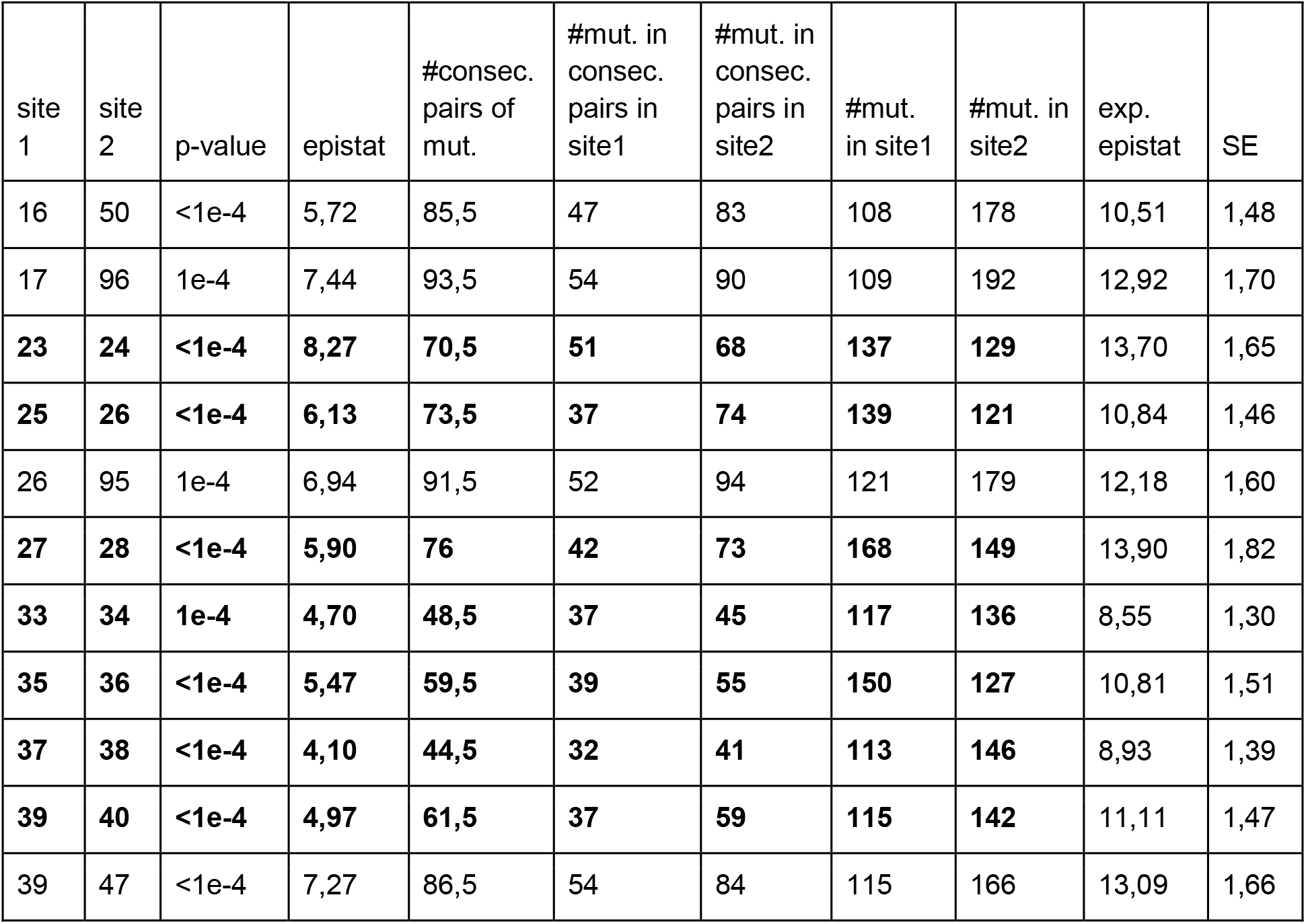
Predicted discordantly evolving pairs for the simulated data with positively and negatively epistatically interacting sites. See legend for table S4. The ‘p-value’ column contains the lower p-values.

**Table S7.**
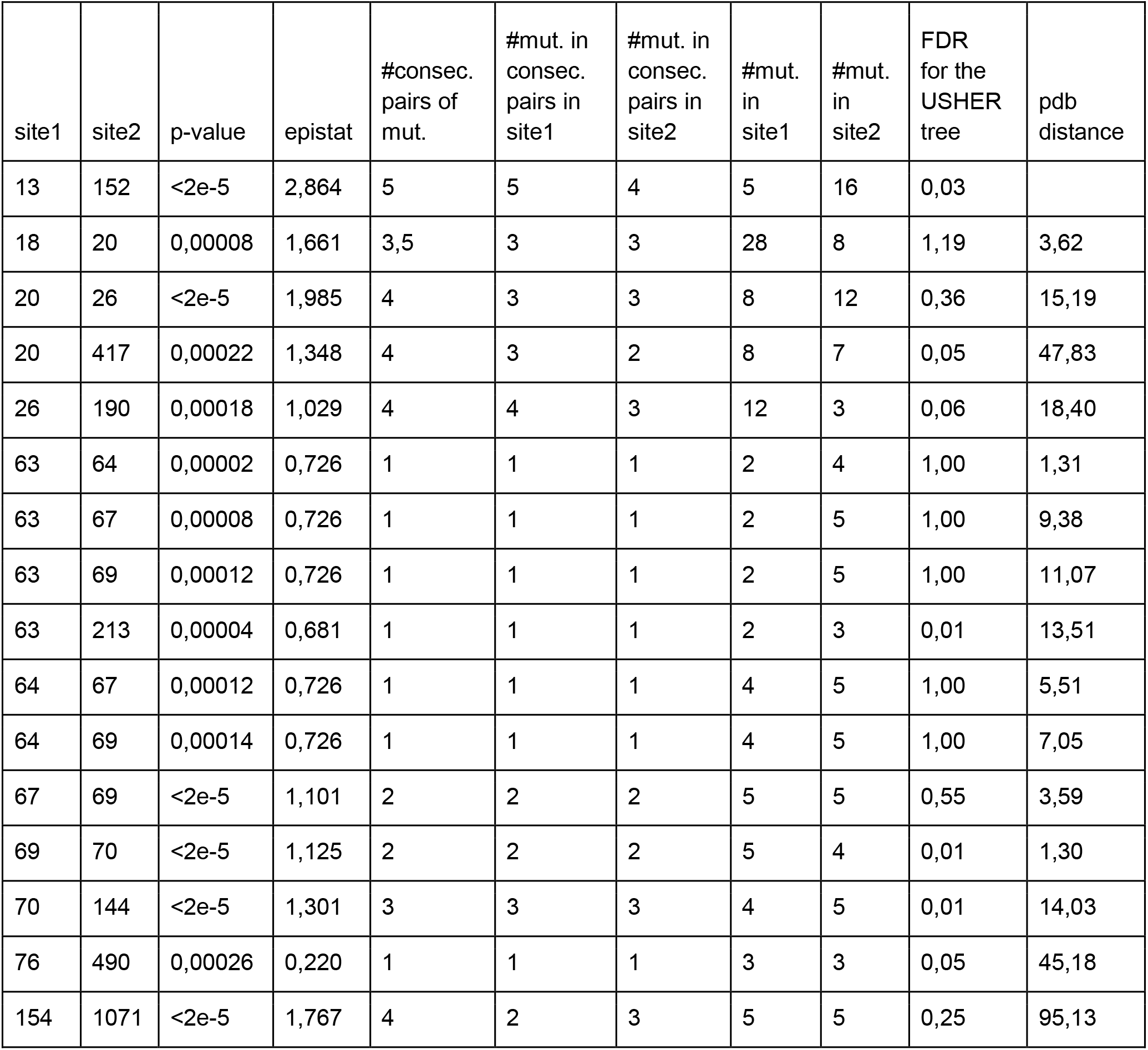
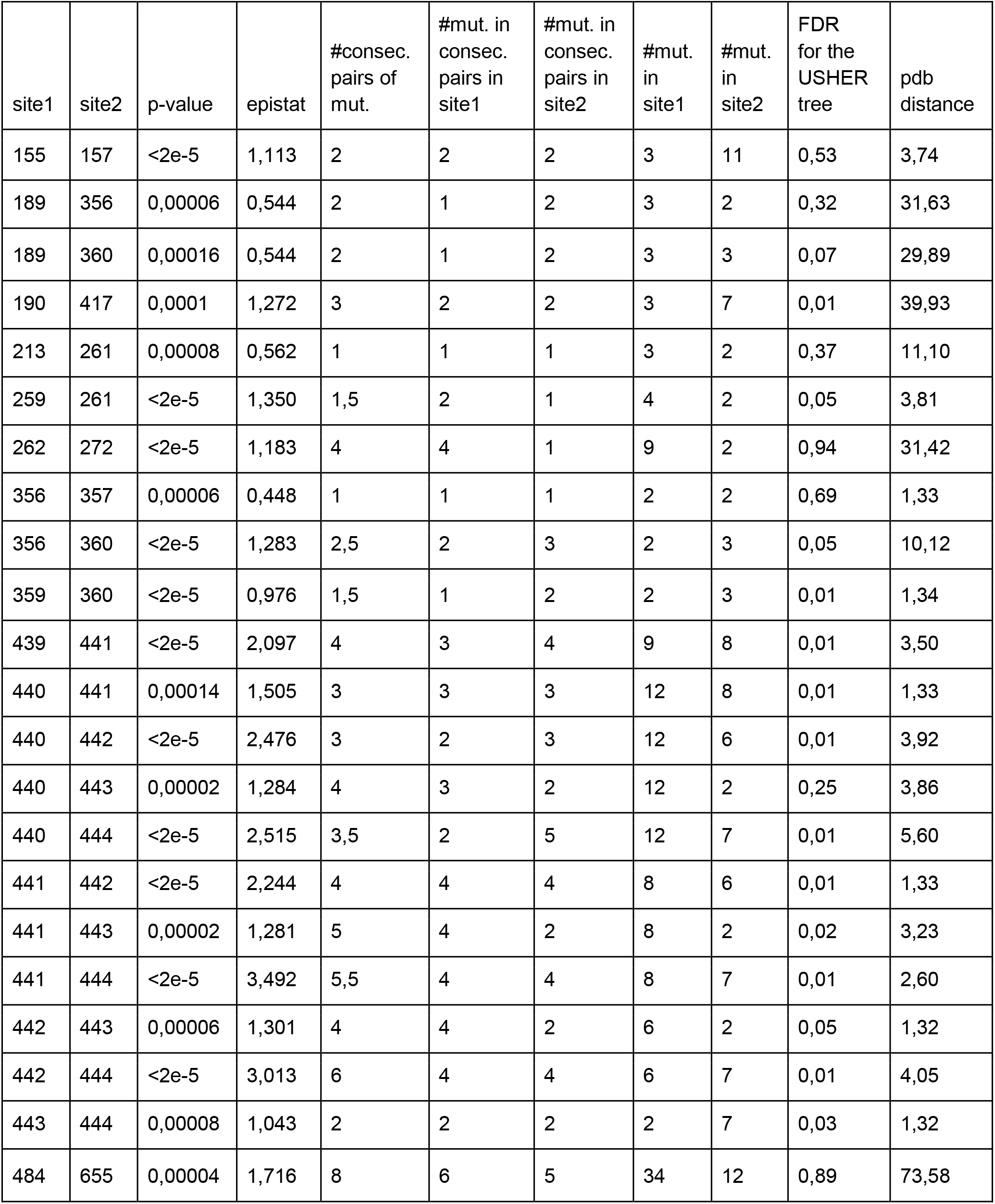
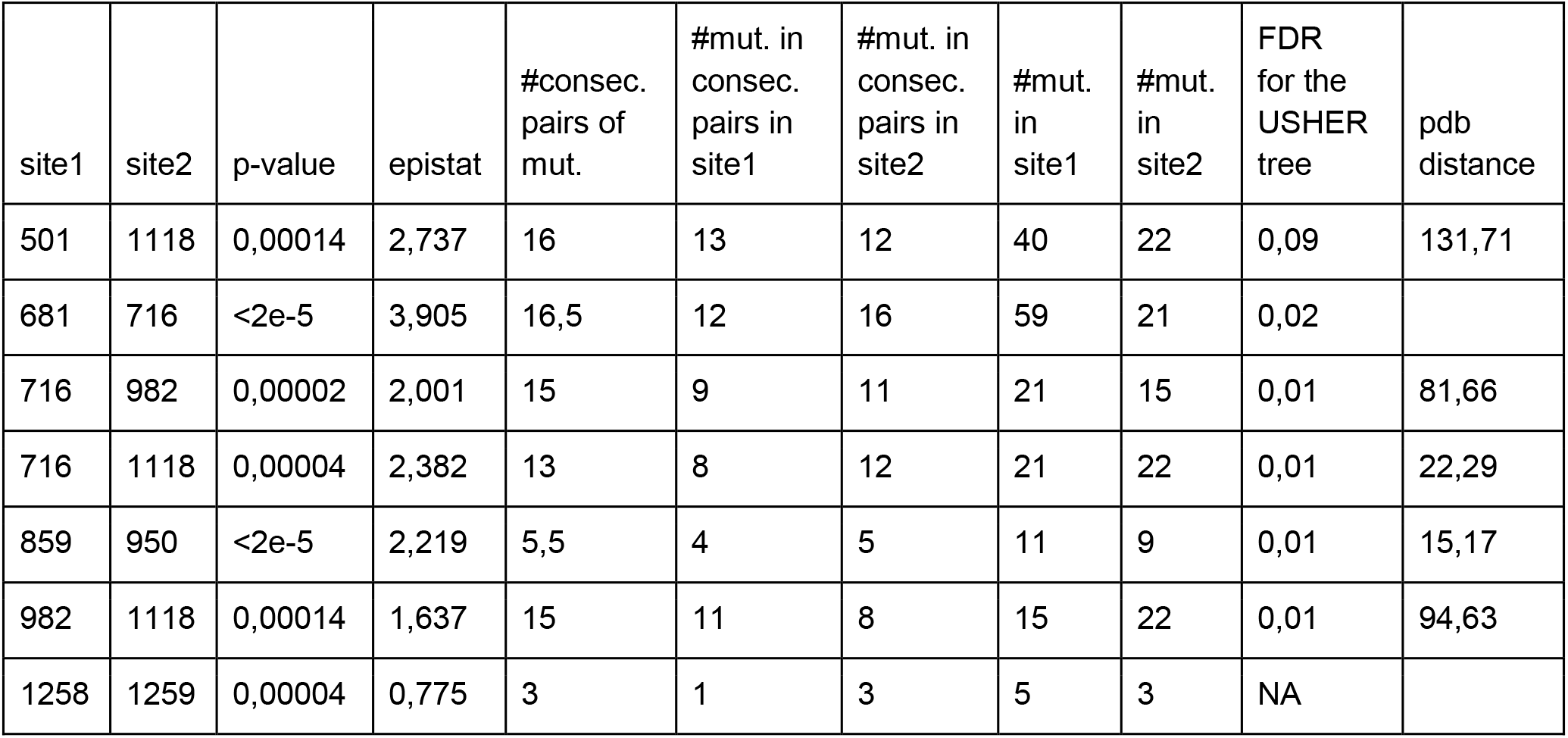
Predicted concordantly evolving pairs for the ML phylogeny of S-gene reconstructed by IQ-TREE. The characteristics of predicted site pairs for the estimated FDR≤10% are shown: the positions of sites on the primary sequence of S-protein (site1 and site2), the upper p-values (p-value), values of the epistatic statistic (epistat), numbers of consecutive pairs of mutations (#consec. pairs of mutations), numbers of mutations in consecutive pairs of sites (#mut. in consec. pairs in site1 and site2) and total numbers of mutations on the internal tree branches in sites (#mut. in site1 and site2), FDR value corresponding to the p-value of the site pair obtained for the alternative phylogeny reconstructed by USHER) and distances between sites on the protein structure 7JJJ (pdb distance)

**Table S8.** Predicted concordantly evolving pairs for the MP phylogeny of S-gene reconstructed by USHER. The characteristics of predicted site pairs for the estimated FDR≤10% are shown: the positions of sites on the primary sequence of S-protein (site1 and site2), the upper p-values (p-value), values of the epistatic statistic (epistat), numbers of consecutive pairs of mutations (#consec. pairs of mutations), numbers of mutations in consecutive pairs of sites (#mut. in consec. pairs in site1 and site2) and total numbers of mutations on the internal tree branches in sites (#mut. in site1 and site2), indicator variable that marks whether a site pairs is also within the set of predictions for the ML phylogeny reconstructed by IQ-TREE for the same FDR threshold (IQ-TREE tree) and distances between sites on the protein structure (pdb distance).

| site1 | site2 | p-value | epistat | #consec. pairs of mut. | #mut. in consec. pairs in site1 | #mut. in consec. pairs in site2 | #mut. in site1 | #mut. in site2 | IQ-TREE | pdb distance |
| --- | --- | --- | --- | --- | --- | --- | --- | --- | --- | --- |
| 18 | 681 | 0,00344 | 0,102 | 5,5 | 5 | 4 | 22 | 62 | 0 |  |
| 222 | 501 | 0,00314 | 0,019 | 1 | 1 | 1 | 12 | 46 | 0 | 55,44 |
| 439 | 501 | 0,00286 | 0 | 0 | 0 | 0 | 10 | 46 | 0 | 5,16 |
| 440 | 501 | 0,0019 | 0 | 0 | 0 | 0 | 12 | 46 | 0 | 9,47 |
| 440 | 681 | 0,00274 | 0,008 | 1 | 1 | 1 | 12 | 62 | 0 |  |
| 484 | 982 | 0,0021 | 0,020 | 3 | 3 | 3 | 43 | 18 | 0 | 35,42 |
| 501 | 675 | 0,00136 | 0,025 | 1 | 1 | 1 | 46 | 22 | 1 | 84,98 |
| 501 | 677 | 0,0001 | 0,023 | 2 | 2 | 2 | 46 | 33 | 1 | 88,2 |
| 570 | 677 | 0,00226 | 0,011 | 1 | 1 | 1 | 19 | 33 | 0 | 44,77 |
| 614 | 653 | 0,00022 | 0,037 | 4 | 1 | 4 | 10 | 5 | 1 | 15,47 |
| 675 | 716 | 0,00568 | 0 | 0 | 0 | 0 | 22 | 21 | 0 | 38,15 |
| 675 | 1118 | 0,00524 | 0 | 0 | 0 | 0 | 22 | 19 | 0 | 58,84 |
| 677 | 681 | 0,00264 | 0,098 | 3 | 3 | 3 | 33 | 62 | 0 |  |
| 677 | 716 | 0,00102 | 0,011 | 1 | 1 | 1 | 33 | 21 | 0 | 42,93 |
| 677 | 982 | 0,00152 | 0,004 | 1 | 1 | 1 | 33 | 18 | 0 | 54,37 |
| 677 | 1118 | 0,00066 | 0,002 | 1 | 1 | 1 | 33 | 19 | 0 | 64,36 |

**Table S9.** Predicted discordantly evolving pairs for the MP phylogeny of S-gene reconstructed by USHER. See legend for the table S8. The ‘p-value’ column contains the lower p-values.

| site1 | site2 | p-value | epistat | #consec.<br>pairs of<br>mut. | #mut. in<br>consec.<br>pairs in<br>site1 | #mut. in<br>consec.<br>pairs in<br>site2 | #mut.<br>in<br>site1 | #mut.<br>in<br>site2 | IQ-TREE | pdb<br>distance |
| --- | --- | --- | --- | --- | --- | --- | --- | --- | --- | --- |
| 12 | 346 | 0,00006 | 0,934 | 2 | 2 | 1 | 9 | 4 | 0 |  |
| 12 | 899 | 0,00004 | 0,724 | 2 | 2 | 1 | 9 | 3 | 0 |  |
| 13 | 152 | 0,00004 | 1,407 | 5 | 4 | 3 | 4 | 13 | 1 |  |
| 20 | 190 | 0,00022 | 0,885 | 3 | 2 | 3 | 8 | 4 | 0 | 22,59 |
| 20 | 417 | 0,00018 | 1,107 | 2 | 2 | 1 | 8 | 8 | 1 | 47,83 |
| 26 | 190 | 0,00026 | 0,913 | 4 | 3 | 4 | 12 | 4 | 1 | 18,40 |
| 26 | 655 | 0,00042 | 0,968 | 6,5 | 3 | 6 | 12 | 14 | 0 | 37,53 |
| 54 | 690 | 0,00044 | 0,714 | 1 | 1 | 1 | 11 | 3 | 0 | 33,65 |
| 62 | 251 | <2e-5 | 0,748 | 1 | 1 | 1 | 2 | 3 | 0 | 32,01 |
| 67 | 96 | <2e-5 | 0,599 | 1 | 1 | 1 | 3 | 7 | 0 | 7,12 |
| 69 | 70 | <2e-5 | 1,496 | 3 | 3 | 3 | 5 | 5 | 1 | 1,30 |
| 69 | 144 | 0,00002 | 0,984 | 2 | 2 | 2 | 5 | 5 | 0 | 12,57 |
| 70 | 144 | <2e-5 | 1,088 | 3 | 3 | 3 | 5 | 5 | 1 | 14,03 |
| 76 | 490 | 0,0001 | 0,275 | 1 | 1 | 1 | 2 | 3 | 1 | 45,18 |
| 80 | 215 | <2e-5 | 1,687 | 6,5 | 3 | 6 | 21 | 14 | 0 | 12,51 |
| 80 | 950 | 0,00014 | 1,091 | 3 | 2 | 2 | 21 | 7 | 0 | 56,59 |
| 152 | 252 | 0,00016 | 0,623 | 2 | 2 | 1 | 13 | 4 | 0 | 15,26 |
| 189 | 360 | 0,0003 | 0,534 | 1 | 1 | 1 | 2 | 2 | 1 | 29,89 |
| 189 | 772 | 0,00024 | 0,440 | 1 | 1 | 1 | 2 | 2 | 0 | 44,17 |
| 190 | 417 | <2e-5 | 1,134 | 2,5 | 2 | 2 | 4 | 8 | 1 | 39,93 |
| 215 | 1167 | 0,00042 | 0,853 | 2 | 2 | 2 | 14 | 4 | 0 |  |

Table S9 cont'd.
| site1 | site2 | p-value | epistat | #consec.<br>pairs of<br>mut. | #mut. in<br>consec.<br>pairs in<br>site1 | #mut. in<br>consec.<br>pairs in<br>site2 | #mut.<br>in<br>site1 | #mut.<br>in<br>site2 | IQ-TREE | pdb<br>distance |
| --- | --- | --- | --- | --- | --- | --- | --- | --- | --- | --- |
| 255 | 256 | 0,00022 | 0,758 | 2 | 2 | 2 | 8 | 7 | 0 | 1,32 |
| 255 | 260 | 0,00036 | 0,811 | 3 | 3 | 2 | 8 | 3 | 0 | 9,61 |
| 256 | 258 | 0,00012 | 0,660 | 1 | 1 | 1 | 7 | 2 | 0 | 2,81 |
| 256 | 260 | 0,00014 | 1,056 | 2 | 2 | 2 | 7 | 3 | 0 | 8,18 |
| 259 | 260 | <2e-5 | 1,536 | 2 | 2 | 2 | 3 | 3 | 0 | 1,31 |
| 259 | 261 | 0,00014 | 0,909 | 1 | 1 | 1 | 3 | 2 | 1 | 3,81 |
| 357 | 360 | <2e-5 | 1,025 | 1,5 | 1 | 2 | 2 | 2 | 0 | 7,59 |
| 359 | 360 | <2e-5 | 1,268 | 2 | 1 | 2 | 2 | 2 | 1 | 1,34 |
| 360 | 772 | 0,00018 | 0,534 | 1 | 1 | 1 | 2 | 2 | 0 | 58,61 |
| 439 | 440 | <2e-5 | 2,038 | 3,5 | 2 | 4 | 10 | 12 | 0 | 1,31 |
| 439 | 441 | <2e-5 | 2,795 | 5 | 3 | 4 | 10 | 5 | 1 | 3,50 |
| 439 | 444 | 0,00006 | 1,606 | 4,5 | 3 | 4 | 10 | 7 | 0 | 6,21 |
| 440 | 441 | <2e-5 | 2,822 | 4,5 | 4 | 2 | 12 | 5 | 1 | 1,33 |
| 440 | 442 | <2e-5 | 2,092 | 3 | 3 | 2 | 12 | 3 | 1 | 3,92 |
| 440 | 444 | <2e-5 | 2,947 | 4,5 | 4 | 3 | 12 | 7 | 1 | 5,60 |
| 441 | 442 | <2e-5 | 2,292 | 3 | 2 | 2 | 5 | 3 | 1 | 1,33 |
| 441 | 443 | 0,00002 | 1,383 | 2 | 2 | 2 | 5 | 2 | 1 | 3,23 |
| 441 | 444 | <2e-5 | 3,829 | 6,5 | 4 | 4 | 5 | 7 | 1 | 2,60 |
| 441 | 445 | 0,00018 | 1,086 | 2 | 2 | 1 | 5 | 6 | 0 | 8,51 |
| 442 | 443 | 0,0001 | 1,132 | 2 | 2 | 1 | 3 | 2 | 1 | 1,32 |

**Table S10.**
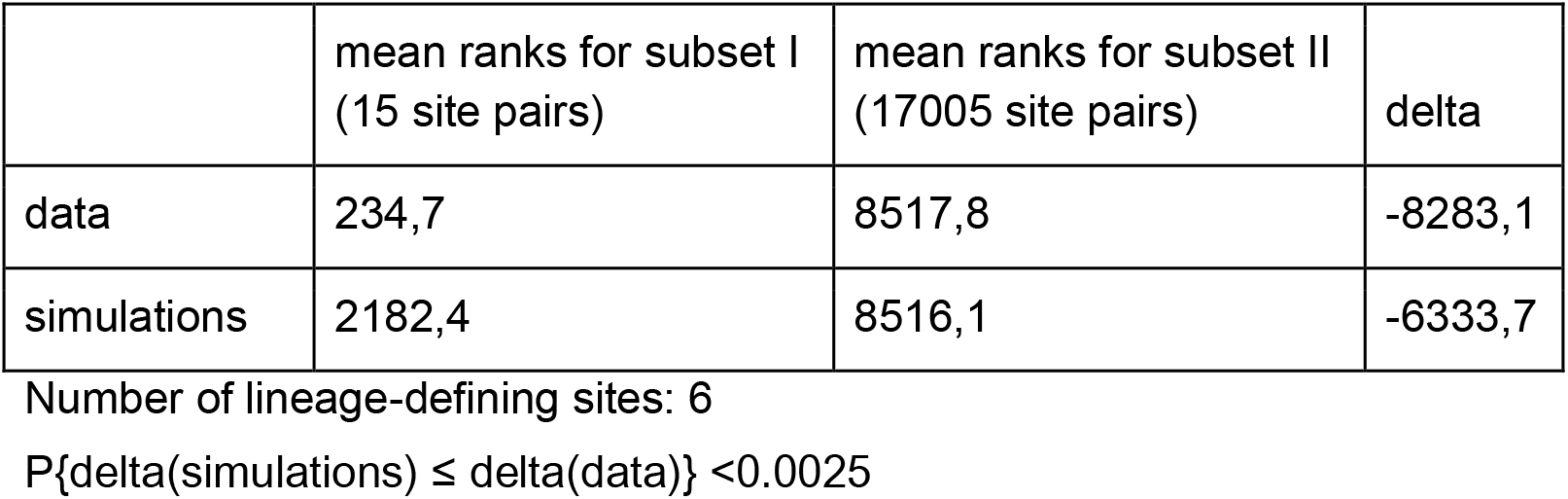
Pairs of lineage-defining sites of Alpha VOC (subset I) tend to have stronger signal of concordant evolution than the complementary subset of site pairs (subset II). Site pairs were ordered by nominal p-values; for each of the two subsets, mean rank of site pairs included into that subset is provided. The difference between mean ranks (delta) obtained for the data is compared to the difference obtained for 400 samples from the null-model where substitutions in each site are independently and randomly distributed on the tree branches (simulations). The probability that the difference of mean ranks for the data is greater than the difference of ranks for samples from the null-model is P{delta(simulations) ≤ delta(data)} <0.0025

**Table S11.** Pairs of lineage-defining sites of Beta VOC tend to have stronger signal of concordant evolution than the complementary subset of site pairs. See legend for the table S10.

|  | mean ranks for subset I<br>(15 site pairs) | mean ranks for subset II<br>(17005 site pairs) | delta |
| --- | --- | --- | --- |
| data | 530,5 | 8517,5 | -7987,1 |
| simulations | 3021,4 | 8515,3 | -5494,0 |

Number of lineage-defining sites: 6
$P\{\text{delta}(\text{simulations}) \leq \text{delta}(\text{data})\} = 0.0025$

**Table S12.** Pairs of lineage-defining sites of Delta VOC tend to have stronger signal of concordant evolution than the complementary subset of site pairs. See legend for the table S10.

|  | mean ranks for subset I<br>(15 site pairs) | mean ranks for subset II<br>(17005 site pairs) | delta |
| --- | --- | --- | --- |
| data | 3137,2 | 8515,2 | -5378,0 |
| simulations | 5545,9 | 8513,1 | -2967,2 |

Number of lineage-defining sites: 6
$P\{\text{delta}(\text{simulations}) \leq \text{delta}(\text{data})\} = 0.005$

**Table S13.** Pairs of lineage-defining sites of Gamma VOC tend to have stronger signal of concordant evolution than the complementary subset of site pairs. See legend for the table S10.

|  | mean ranks for subset I<br>(55 site pairs) | mean ranks for subset II<br>(16965 site pairs) | delta |
| --- | --- | --- | --- |
| data | 543,3 | 8536,3 | -7993,0 |
| simulations | 3532,3 | 8526,6 | -4994,3 |

Number of lineage-defining sites: 11

**Table S14.** Pairs of lineage-defining sites of Omicron VOC tend to have stronger signal of concordant evolution than the complementary subset of site pairs. See legend for the table S10.

|  | mean ranks for subset I<br>(91 site pairs) | mean ranks for subset II<br>(16929 site pairs) | delta |
| --- | --- | --- | --- |
| data | 5686,0 | 8525,7 | -2839,7 |
| simulations | 5540,6 | 8526,5 | -2985,9 |

Number of lineage-defining sites: 14

**Table S15.** Pairs of lineage-defining sites of Alpha VOC tend to have stronger signal of concordant evolution than the complementary subset of site pairs even if all Alpha and related lineages are excluded from the analysis. See legend for the table S10.

|  | mean ranks for subset I<br>(15 site pairs) | mean ranks for subset II<br>(15385 site pairs) | delta |
| --- | --- | --- | --- |
| data | 316,8 | 7707,7 | -7390,9 |
| simulations | 4281,3 | 7703,8 | -3422,5 |

Number of lineage-defining sites: 6

**Table S16.** No difference in the strength of concordant evolution is detected for pairs of lineage-defining sites of Beta VOC and the complementary subset of site pairs if all Beta and related lineages are excluded from the analysis. See legend for the table S10.

|  | mean ranks for subset I<br>(15 site pairs) | mean ranks for subset II<br>(16275 site pairs) | delta |
| --- | --- | --- | --- |
| data | 4586,7 | 8148,8 | -3562,1 |
| simulations | 4522,3 | 8148,8 | -3626,6 |

Number of lineage-defining sites: 6

**Table S17.** No difference in the strength of concordant evolution is detected for pairs of lineage-defining sites of Delta VOC and the complementary subset of site pairs if all Delta and related lineages are excluded from the analysis. See legend for the table S10.

|  | mean ranks for subset I<br>(15 site pairs) | mean ranks for subset II<br>(16638 site pairs) | delta |
| --- | --- | --- | --- |
| data | 5934,7 | 8329,2 | -2394,5 |
| simulations | 5975,1 | 8329,1 | -2354,0 |

Number of lineage-defining sites: 6

**Table S18.** Pairs of lineage defining sites of Gamma tend to have weaker signal of concordant evolution than the complementary subset of site pairs if all Gamma and related lineages are excluded from the analysis. See legend for the table S10.

|  | mean ranks for subset I<br>(45 site pairs) | mean ranks for subset II<br>(16245 site pairs) | delta |
| --- | --- | --- | --- |
| data | 6424,4 | 8150,3 | -1725,8 |
| simulations | 5393,5 | 8153,1 | -2759,7 |

Number of lineage-defining sites: 10

**figure S1.**
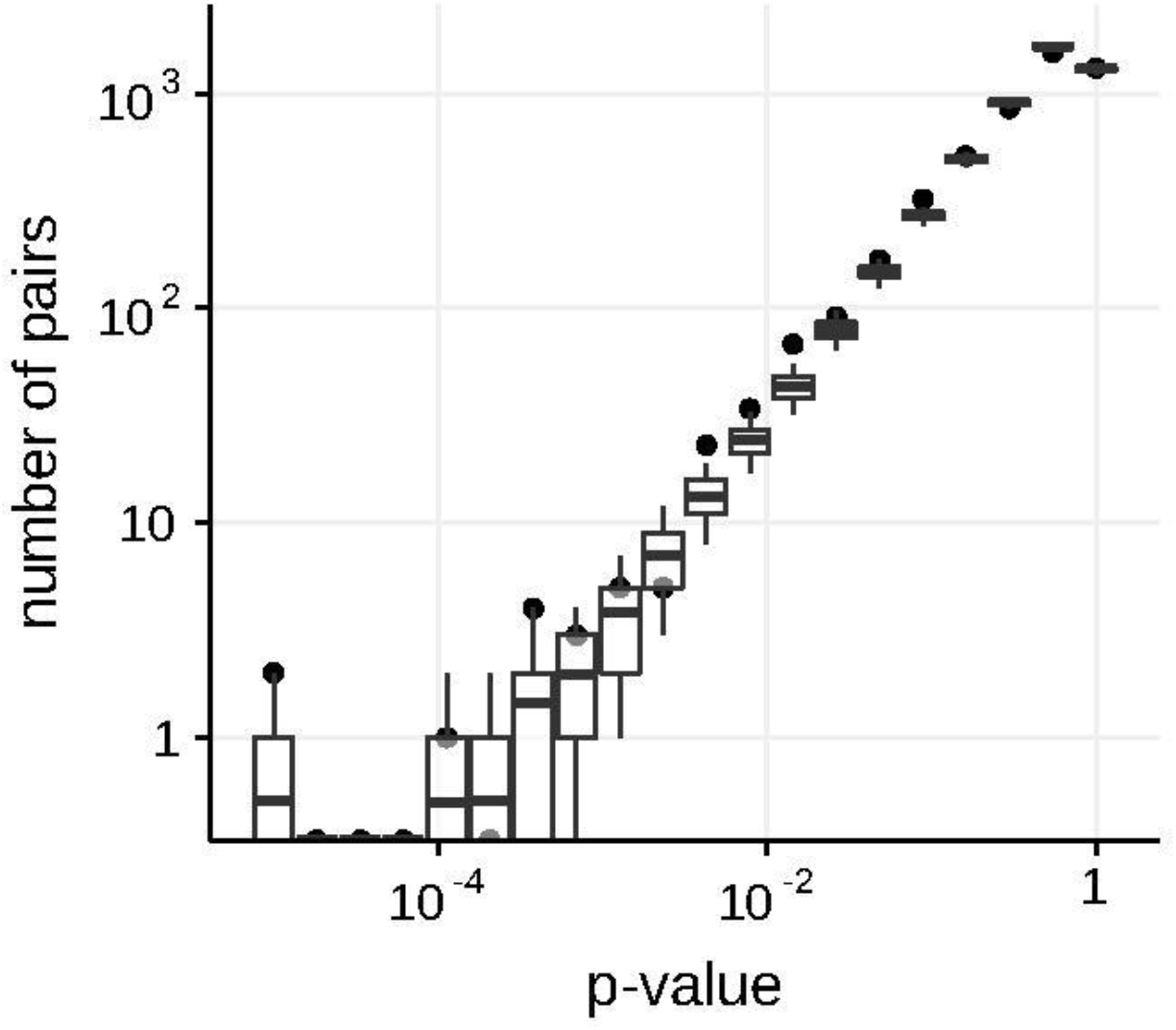
Numbers of predicted concordantly evolving pairs for different nominal p-values in the simulated data in non-epistatic mode of evolution, compared to the null distribution for the ML phylogeny reconstructed by IQ-TREE. Black dots indicate data points, boxes with whiskers indicate simulation results. Top and bottom of each box correspond to the 75th and 25th percentile, whiskers correspond to the 95th and 5th percentile. Vertical line corresponds to the 10% FDR threshold.

**Figure S2.**
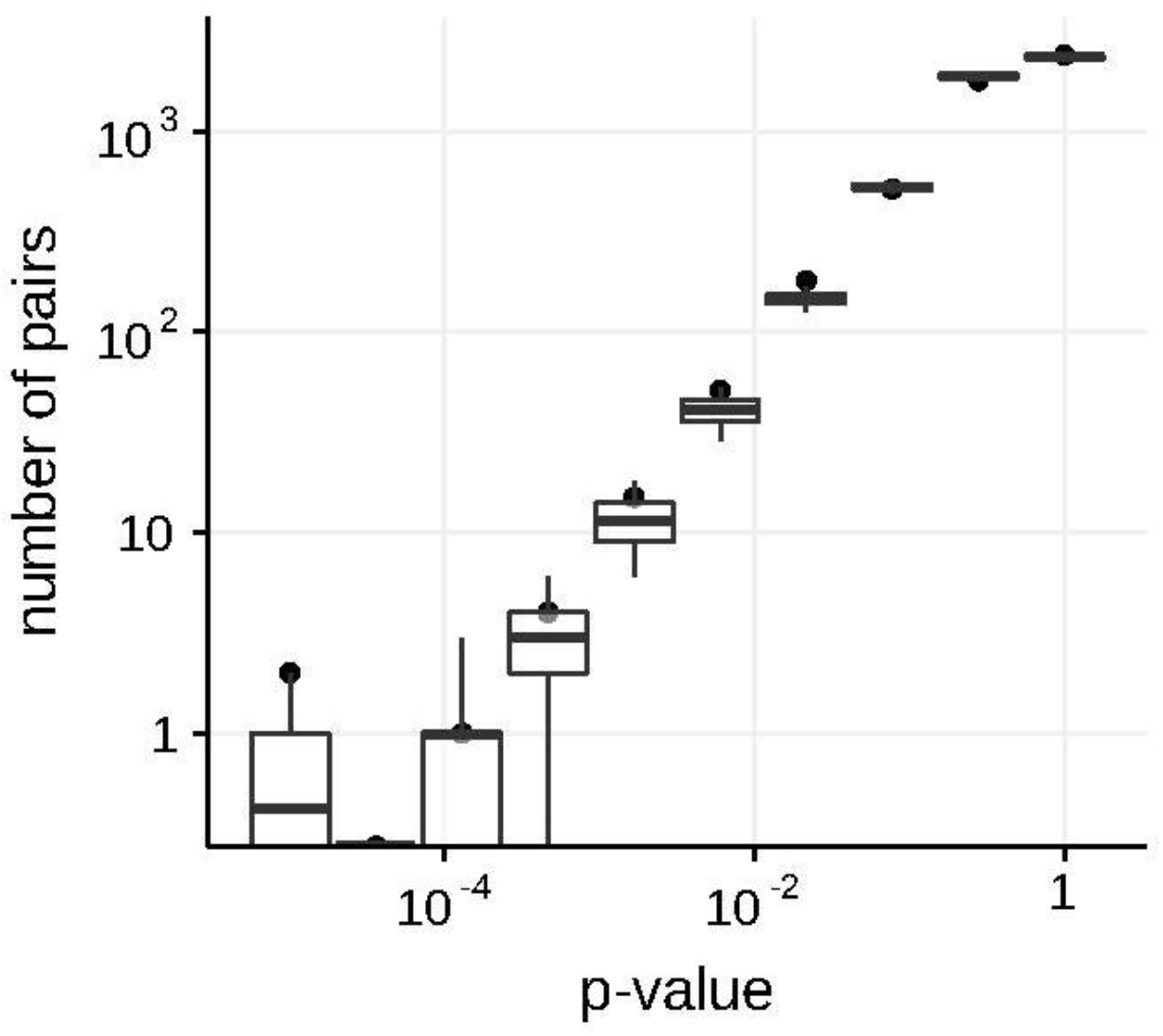
Numbers of predicted discordantly evolving pairs for different nominal p-values in the simulated data in non-epistatic mode of evolution, compared to the null distribution for the ML phylogeny reconstructed by IQ-TREE. See legend for the Fig. S1.

**Figure S3.**
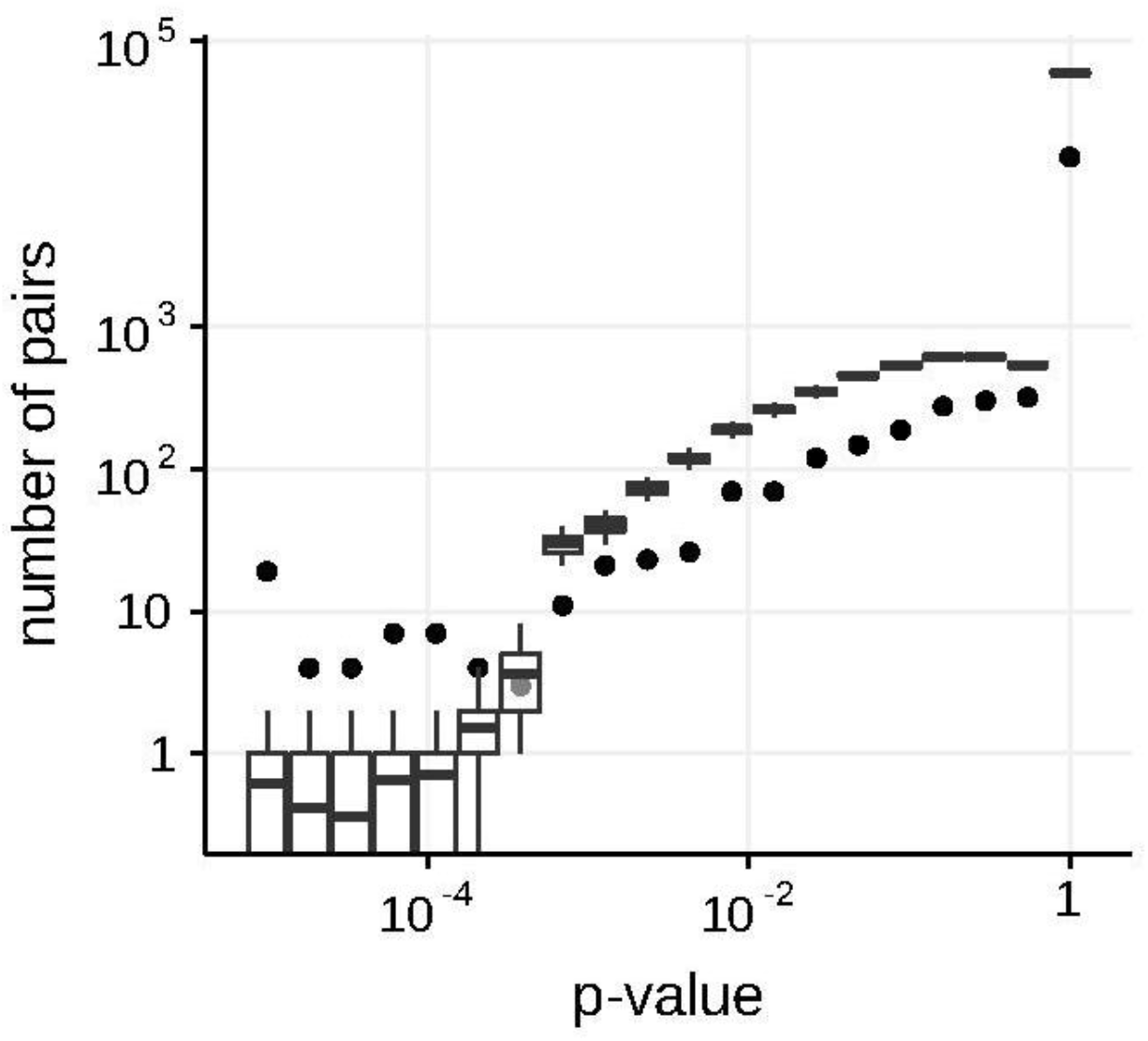
Numbers of predicted concordantly evolving pairs for different nominal p-values in the S-gene of SARS-Cov-2, compared to the null distribution for the ML phylogeny reconstructed by IQ-TREE. See legend for the Fig. S1.

**Figure S4.**
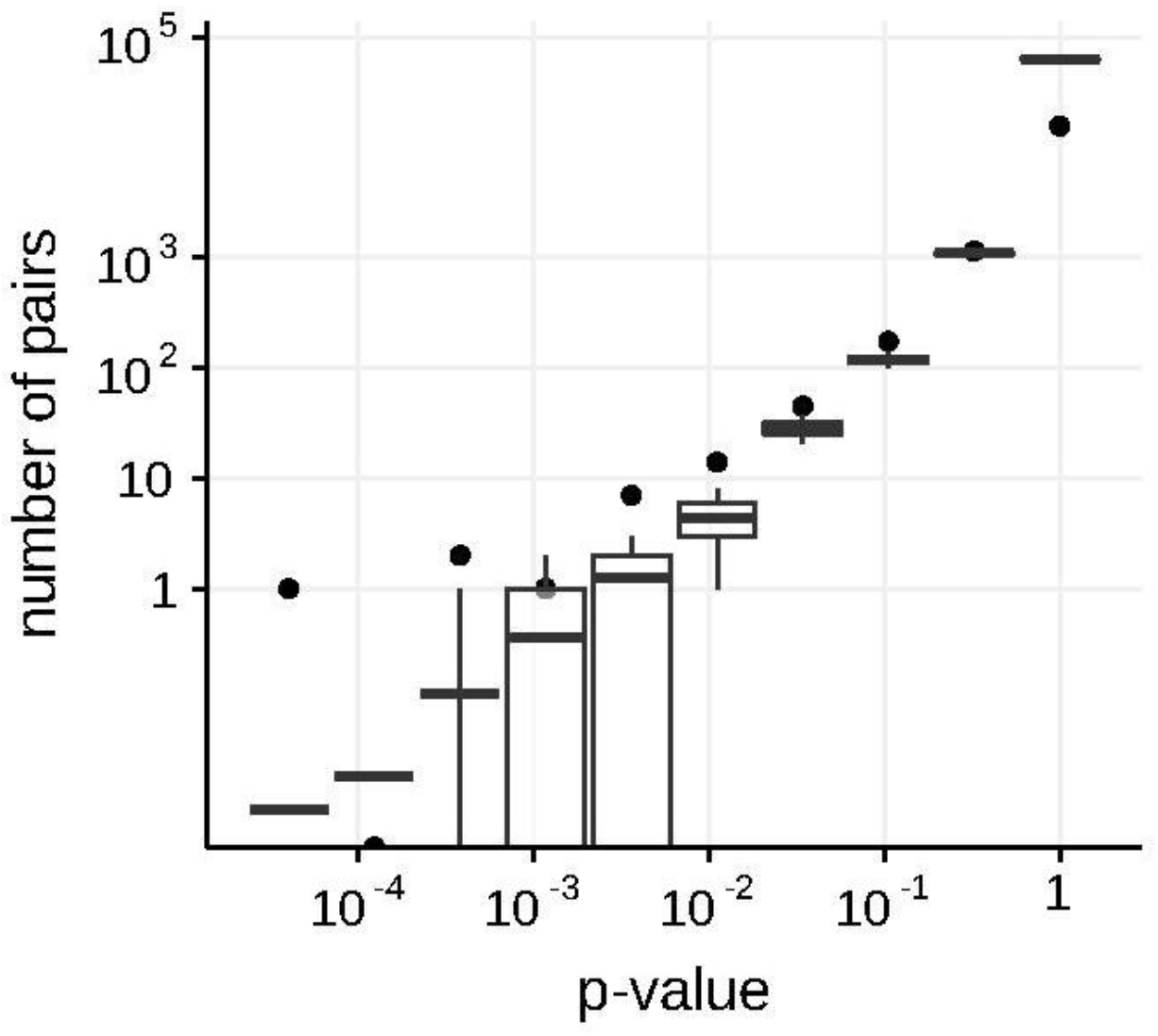
Numbers of predicted discordantly evolving pairs for different nominal p-values in the S gene of SARS-Cov-2, compared to the null distribution for the ML phylogeny reconstructed by IQ-TREE. See legend for the Fig. S1.

